# Mouse dLGN receives input from a diverse population of retinal ganglion cells with limited convergence

**DOI:** 10.1101/322164

**Authors:** Miroslav Román Rosón, Yannik Bauer, Philipp Berens, Thomas Euler, Laura Busse

**Affiliations:** Centre for Integrative Neuroscience, University of Tübingen, Germany.; Institute for Ophthalmic Research, University Hospital Tübingen, Germany.; Division of Neurobiology, Department Biology II, LMU Munich, Germany; Graduate School of Neural & Behavioural Sciences | International Max Planck Research School, University of Tübingen, Germany.; Graduate School of Systemic Neuroscience (GSN), LMU Munich, Germany; Bernstein Centre for Computational Neuroscience, Germany

## Abstract

In the mouse, the parallel output of more than 30 functional types of retinal ganglion cells (RGCs) serves as the basis for all further visual processing. Little is known about how the representation of visual information changes between the retina and the dorsolateral geniculate nucleus (dLGN) of the thalamus, the main relay station between the retina and cortex. Here, we functionally characterized responses of retrogradely labeled dLGN-projecting RGCs and dLGN neurons to the same set of visual stimuli. We found that many of the previously identified functional RGC types innervate the dLGN, which maintained a high degree of functional diversity. Using a linear model to assess functional connectivity between RGC types and dLGN neurons, we found that the responses of dLGN neurons could be predicted as a linear combination of inputs from on average five RGC types, but only two of those had the strongest functional impact. Thus, mouse dLGN receives input from a diverse population of RGCs with limited functional convergence.

## INTRODUCTION

In the mammalian retina, the photoreceptor signal is decomposed into multiple parallel channels ^1,2^, carried to the brain by more than 30 types of retinal ganglion cells (RGCs) ^3,4^. Each type of RGC extracts specific features of the visual environment, which are projected via the optic nerve to more than 50 retino-recipient areas in the brain ^5,6^. Among those, a key center transmitting information to the primary visual cortex (V1) is the dorsolateral geniculate nucleus (dLGN) of the thalamus ^7^.

Retino-geniculate information transmission has been studied extensively in cats and monkeys, where the vast majority of dLGN neurons seems to be driven by only few (1–3) dominant RGCs ^8,9^. This dominant input can evoke such strong excitatory postsynaptic potentials (EPSPs) - so-called “S-potentials” - that they can be picked up by extracellular recordings. Consistent with a low degree of convergence, the S-potentials of a dLGN neuron and its spiking output have closely matching receptive fields (RFs) in terms of location, center-surround organization and size ^10–12^. In addition to these dominant inputs, cat dLGN cells receive weaker input from multiple RGCs as shown by electron microscopy^13^ and *in vivo* electrophysiology^8,9,14,15^. Given the strict spatial layering of the dLGN in these species, it is generally assumed that the inputs into dLGN cells arise from RGCs of the same type, although at least 13 types of dLGN-projecting RGCs have been identified in monkeys ^16^.

Recent anatomical studies have started to explore retino-geniculate connectivity in the mouse, revealing a complex picture. Mono-transsynaptic rabies virus tracing of inputs received by individual neurons in dLGN demonstrated that mouse dLGN neurons can be divided into two groups based on the pattern of their retinal inputs ^17^: while some dLGN cells received inputs from mostly a single RGC type (“relay input mode”), others showed a high degree of convergence, with inputs being composed of up to several dozen RGCs of different types (“combination input mode”). A high degree of retino-geniculate convergence is further supported by recent ultrastructural studies of retinal afferents and their thalamic relay cell targets ^18,19^.

Given the functional diversity of RGCs ^20^ and the complex input patterns of dLGN neurons ^21^, one would expect an at least similarly rich functional representation in mouse dLGN. In contrast, the majority of mouse dLGN neurons has been reported to have circularly symmetric RFs and is believed to perform linear spatial summation ^22,23^, similar to results in primates ^24^. However, mouse dLGN also contains neurons with more complex and diverse response properties: orientation-selective (OS) and direction-selective (DS) cells tuned along the four cardinal axes ^25–27^, as well as a significant number of “suppressed-by-contrast” cells signaling uniformity of the visual field ^26^. Finally, a heterogeneous population of cells with long latencies and responses to both the on- and offset of light has been reported ^26^. It is currently unknown whether these response properties are inherited from the innervating retinal afferents or emerge de-novo in dLGN by a combination of converging retinal inputs and dLGN-intrinsic computations.

Here, we sought to determine how the visual representation in mouse dLGN arises from retinal inputs. We show that many previously identified functional RGC types ^3^ innervate the dLGN, which yields diverse geniculate representations. These dLGN responses can be modelled by a linear combination of on average five RGC types, amongst which two have the strongest functional impact. We conclude that mouse dLGN neurons receive input from multiple distinct retinal output channels and relay diverse retinal representations of visual information to the cortex, with limited functional convergence.

## RESULTS

### Retrograde viral tracing to functionally characterize dLGN-projecting RGCs

To identify dLGN-projecting (dLGN-p) RGCs, we injected a Cre-encoding retrograde Herpes-Simplex-Virus 1 (LT HSV-hEF1a-cre)^28^ into the dLGN of a transgenic reporter mouse line with a floxed genetically encoded Ca^2+^ indicator (GCaMP6f) ^29,30^. After transducing the axon terminals ^31,32^ in the dLGN, the virus was retrogradely transported to the soma, where it triggered the expression of Cre-recombinase and, subsequently, the Cre-dependent expression of GCaMP6f. Since the virus does not spread trans-synaptically, it only labeled cells with afferents in the dLGN. This enabled us to label only the subset of dLGN-p RGCs in the retina (Fig. 1a).

**Figure 1.**
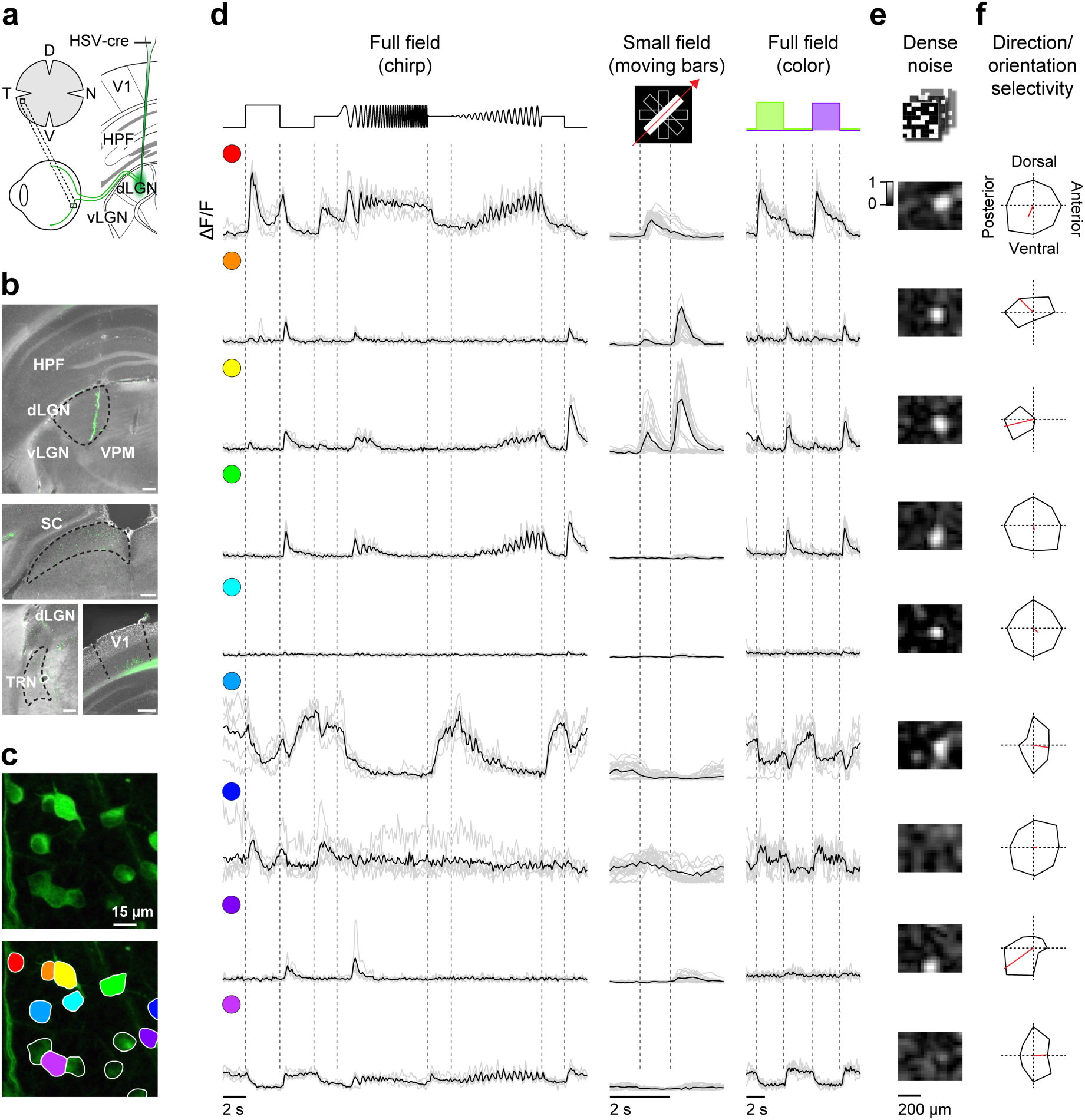
| Functional characterization of dLGN-projecting RGCs. **a,** Schematic of the experimental approach. **b,** Injection of a Cre-encoding retrograde Herpes Simplex Virus 1 (LT HSV-hEFIα-cre)^28^ into the dLGN (green, GCaMP6f; grey DAPI). Injection site (top) and areas with retrogradely labeled cell bodies (below) and outlines of the dorsolateral geniculate complex (dLGN), superior colliculus (SC), reticular nucleus of the thalamus (TRN) and primary visual cortex (V1). Scale bar: 200 μm. **c,** Whole-mounted mouse retina of a floxed GCaMP6f mouse transduced with LT HSV-hEFIα-cre and recorded with a two-photon microscope in the ganglion cell layer. Top: Scan field (110 × 110 μm). Bottom: Regions of interest (ROIs) on GCaMP6f expressing RGCs. **d,** Ca^2+^ responses ∆F/F) from 9 exemplary ROIs color-coded in (c) and evoked by three visual stimuli: Full-field chirp, bright bars moving in eight directions, and full-field alternating green/blue. Single trials in grey, averages of n = 5 (chirp), 7 (green/blue) or 24 (moving bars) trials in black. Traces are scaled to the maximal value of ΔF/F for each stimulus separately over all ROIs to make ROI intensities comparable. **e**,**f,** spatial RFs (e) and polar plots indicating direction and orientation selectivity (f; vector sum in red) for the same 9 cells as in (d). HPF, hippocampus; vLGN, ventral part of the lateral geniculate complex; VPM, ventral posteromedial nucleus of the thalamus.

We histologically confirmed the target location of the virus injection, as well as its limited diffusion. In line with earlier studies, we found retrogradely labeled neurons in additional dLGN-projecting structures, including the superior colliculus (Fig. 1b, center), the thalamic reticular nucleus and the deep layers of primary visual cortex (Fig. 1b, bottom) ^33,34^.

We then used two-photon Ca^2+^ imaging to measure the light-evoked responses of the dLGN-p RGCs (Fig. 1c-f). We probed their response properties across the whole retina with a standardized stimulus set used in a previous RGC classification study ^3^. The Ca^2+^-responses of the labeled RGCs (Fig. 1c, top) were analyzed using manually drawn regions of interest (ROIs) (Fig. 1c, bottom). In total, we identified 581 virus-labeled RGCs as ROIs, with a range of 1–25 ROIs per field (median = 6). In addition to the classical ON/OFF-response types and the direction selective RGCs ^35,36^, we found cells among the dLGN-p RGCs that, for example, responded differently to local and full-field stimuli, showed preference to higher or lower frequency stimulation or were suppressed by frequency and contrast (Fig. 1d-f).

### The majority of functional RGC types project to dLGN

We next asked which of the previously characterized functional mouse RGC types ^3^ project to the dLGN. To this end, we used the functional RGC clusters obtained previously (henceforth “RGC-all clusters”), and identified, for each retrogradely labeled dLGN-p RGC, the RGC-all cluster with the best matching response properties (Supp. Fig. 1). To account for the differences in Ca^2+^ indicators between the two studies, we first de-convolved the Ca^2+^ signals of the GCaMP6f data set and the published OGB-1 data set ^3^ using the respective Ca^2+^ kernels for the two indicators (Supp. Fig. 2). We then correlated, for both the "chirp” stimulus and the moving bar stimulus, the trial-averaged responses of each dLGN-p RGC to the mean response of each RGC-all cluster and combined the correlation coefficients weighted by a stimulus-specific response quality index (Qi) into a “match index” (Mi). We assigned each dLGN-p RGC to the RGC-all cluster with highest Mi. For both individual example cells (Fig. 2a) and across the population of dLGN-p RGCs, the assignment worked well (median Mi = 0.62). Accordingly, population mean responses of dLGN-p RGCs assigned to the same cluster agreed well with the population mean responses of their RGC-all cluster (Fig. 2b).

**Figure 2.**
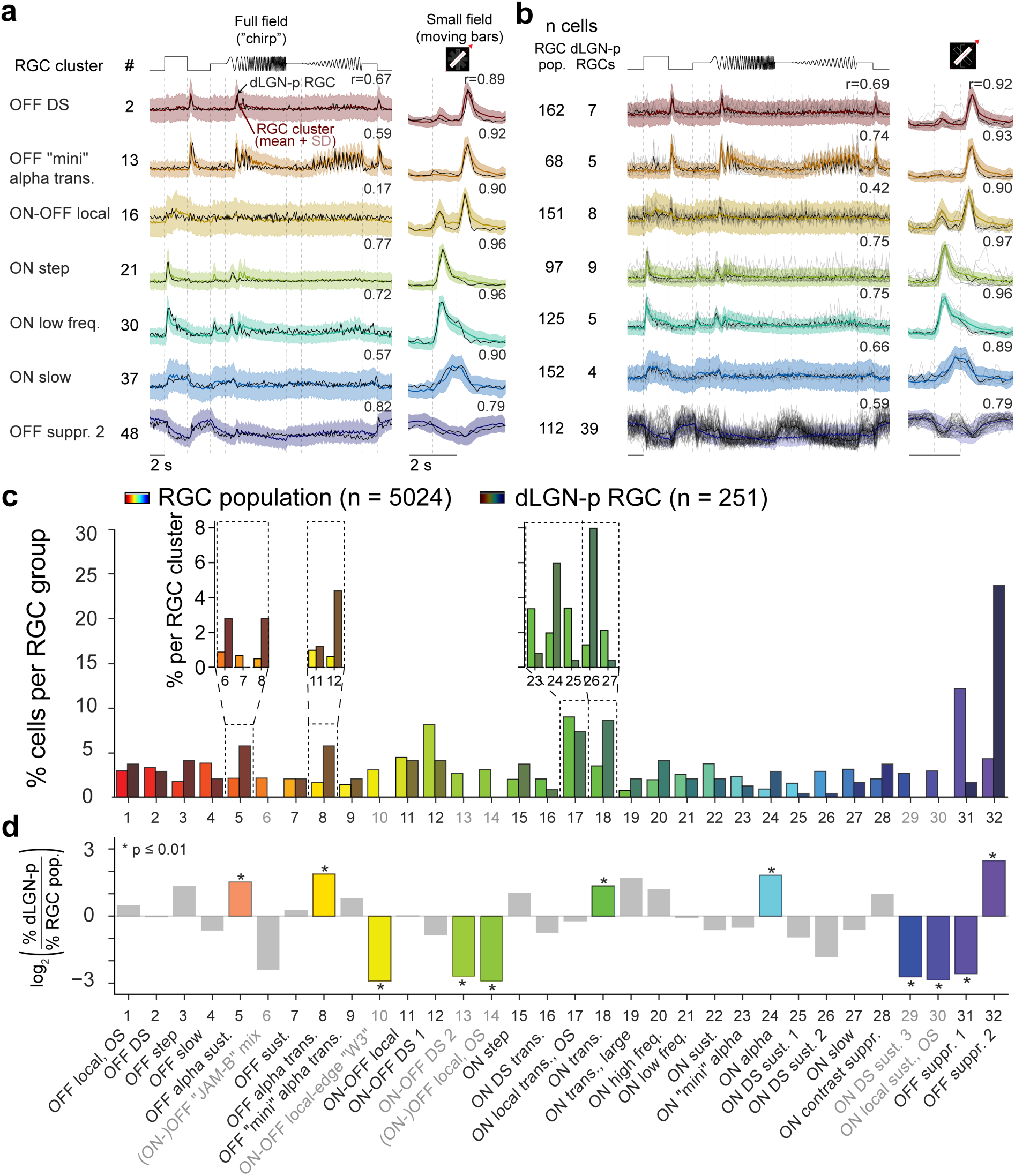
| Cluster assignments. **a,** Responses of selected RGC-all clusters (colored line, mean; shaded area, ±1 SD) with mean responses of assigned example dLGN-p RGCs (black) to chirp (left) and bar stimulus (right). Numbers indicate correlation coefficients. **b,** Same, for all assigned dLGN-p RGCs in the corresponding clusters (grey, responses of single dLGN-p RGCs; black, mean). **c,** Distribution of cells per RGC-all group from ref ^3^ vs. cells per dLGN-p group; insets illustrate subdivision of selected groups into contributing clusters. **d**, Comparison of cell-per-group percentages as log_2_-ratio (%dLGN-p RGCs / %RGC-all). Significant differences in cell proportions (p < 0.01; binomial test) are marked as colored bars and with asterisks.

We then determined which RGC types were over- or underrepresented in the dLGN-p RGC population compared to the complete RGC population. Following ref ^3^, we grouped the 49 RGC-all clusters into 32 groups, where each group represents an RGC type based on knowledge not only about functional features, but also about soma size, receptive field (RF) size, and direction and orientation selectivity (Fig. 2c, for details see Methods; see Supp. Fig. 3 for same analysis without grouping). Surprisingly, 75% of RGC-all groups were assigned dLGN-p RGCs (24/32 groups with at least two cells), suggesting that the majority of RGC types projects to the dLGN.

Several RGC types were systematically over- or underrepresented in the dLGN-p population (Fig. 2c,d; binomial test, P<0.01): Overrepresented were “OFF-suppressed 2” cells (Fig. 2c,d: group (G) 32 / Fig. 2c inset: clusters (C) 47–49) and all classical alpha cells (Fig. 2c,d), including OFF alpha sust. (G_5_ / C_6,7,8_), OFF alpha trans. (G_8_ / C_11,12_), and ON alpha (G_24_ / C_34_), as well an ON transient RGC (G_18_ / C_34_). Underrepresented, in turn, were the ON-OFF local-edge “W3” (G_10_ / C_14_), and several orientation- and direction-selective RGCs (i.e., ON-OFF DS 2 (G_13_ / C_19_), (ON-)OFF local OS (G_14_ / C_20_), ON DS sust. 3 (G_20_ / C_40_) and ON local sust. OS (G_30_ / C_41_). Other groups contributed roughly the same percentage to the dLGN-p cells as to the total RGC population (for details, see Discussion). These results indicate that dLGN receives diverse, parallel input from many functional RGC groups, with a striking overrepresentation of alpha and of one group of suppressed-by-contrast RGCs.

### The dLGN population response is composed of diverse components

Having established that many functional RGC types provide input to mouse dLGN, we next wondered how this diversity is reflected in the dLGN population. To study this question, we performed extracellular single-unit recordings of geniculate neurons in head-fixed mice (Fig. 3a; Supp. Fig. 1). We verified the recording sites by histological reconstruction of the electrode tract (Fig. 3b) and the characteristic progression of retinotopy along the electrode channels (Fig. 3c, see also ref ^26^). We presented the same full-field chirp stimulus as in the retina experiments. To assess the stability of the dLGN recordings and the consistency of our spike sorting, we flanked the chirp stimulus by presentations of drifting gratings with varying orientation, temporal and spatial frequency.

**Figure 3.**
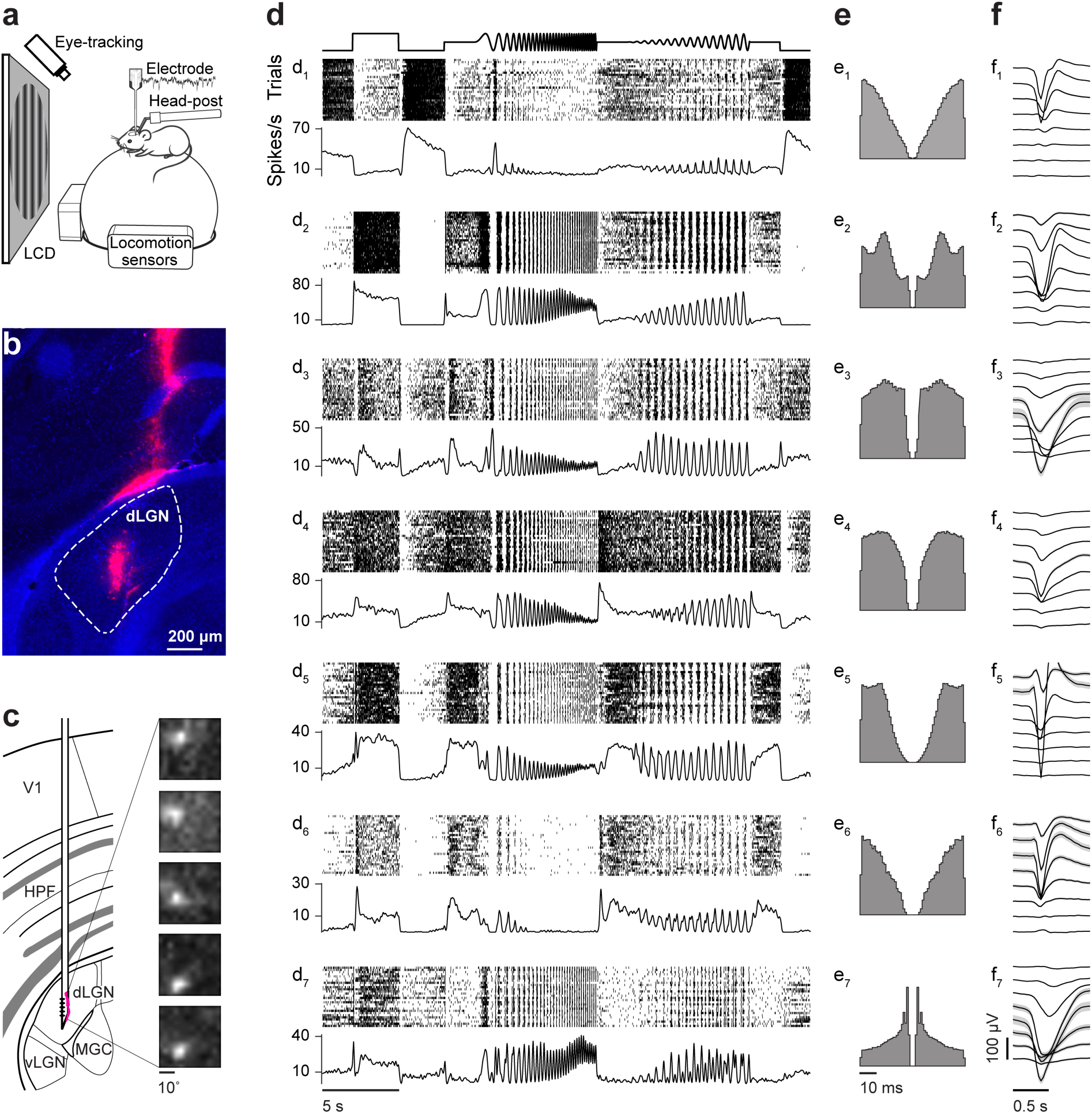
| Functional characterization of dLGN neurons. **a**, Schematic of the experimental setup for extracellular dLGN recordings. **b,** Reconstruction of the electrode track in dLGN (coronal section; dLGN is marked by white outline, Blue: DAPI, Red: DiD coating the electrode). **c**, Schematic of recording site with RFs mapped for several electrode channels (dorsal to ventral), showing the retinotopic progression in elevation typical of mouse dLGN. **d-f**, Spike raster plots (top) and spike density function (bottom) of 7 exemplary dLGN neurons in response to the chirp stimulus (d), their autocorrelograms (e), and spike waveforms in 5 selected channels of the 32-channel probe (f). V1, primary visual cortex; HPF, hippocampus; dLGN, dorsal part of the lateral geniculate nucleus; vLGN, ventral part of the lateral geniculate nucleus; MGC, medial geniculate complex.

Responses of dLGN neurons to the chirp stimulus were surprisingly diverse: The cells not only displayed the “standard” transient and sustained ON/OFF responses, or were suppressed by contrast, but also differed in their temporal frequency or contrast preferences as well as their response kinetics (Fig. 3d). Some of the cells even displayed slow ramping responses (Fig. 3d_4_, d_5_). To ensure that this response diversity did not result from poor unit isolation during spike sorting, we considered only units with a firing rate > 1 spike/s, which were stable across time, had a clean refractory period and a distinct extracellular spike waveform (Fig. 3e, f). This diversity is likely also not related to influences of locomotion (Supp. Fig. 5).

To quantitatively assess the degree of diversity present in the dLGN neuron population, we decomposed the single unit responses to the chirp stimulus into “response components” using non-negative matrix factorization (NNMF) ^37^. Individual neuron responses can then be reconstructed as a weighted sum of these elementary response components (Fig. 4a, Supp. Fig. 1). Interestingly, decomposing our dLGN population response into only two fundamental components revealed components resembling responses with ON and OFF features; increasing the rank number to four, added temporal diversity with transient and sustained features; increasing rank number further led to additional contrast-suppressed features (Fig. 4b. To determine the mathematically optimal number of components, we used cross-validation^38^ (Fig. 4c), where we repeatedly performed the factorization on different training sets for an increasing number of components (ranks) (Fig. 4b), and evaluated the mean squared errors (MSEs) of the NNMF model on validation sets not used for factorization (for details, see Methods). This procedure yielded an optimal number of components at k = 33 (Fig. 4d,e), indicative of a large diversity of dLGN responses.

**Figure 4.**
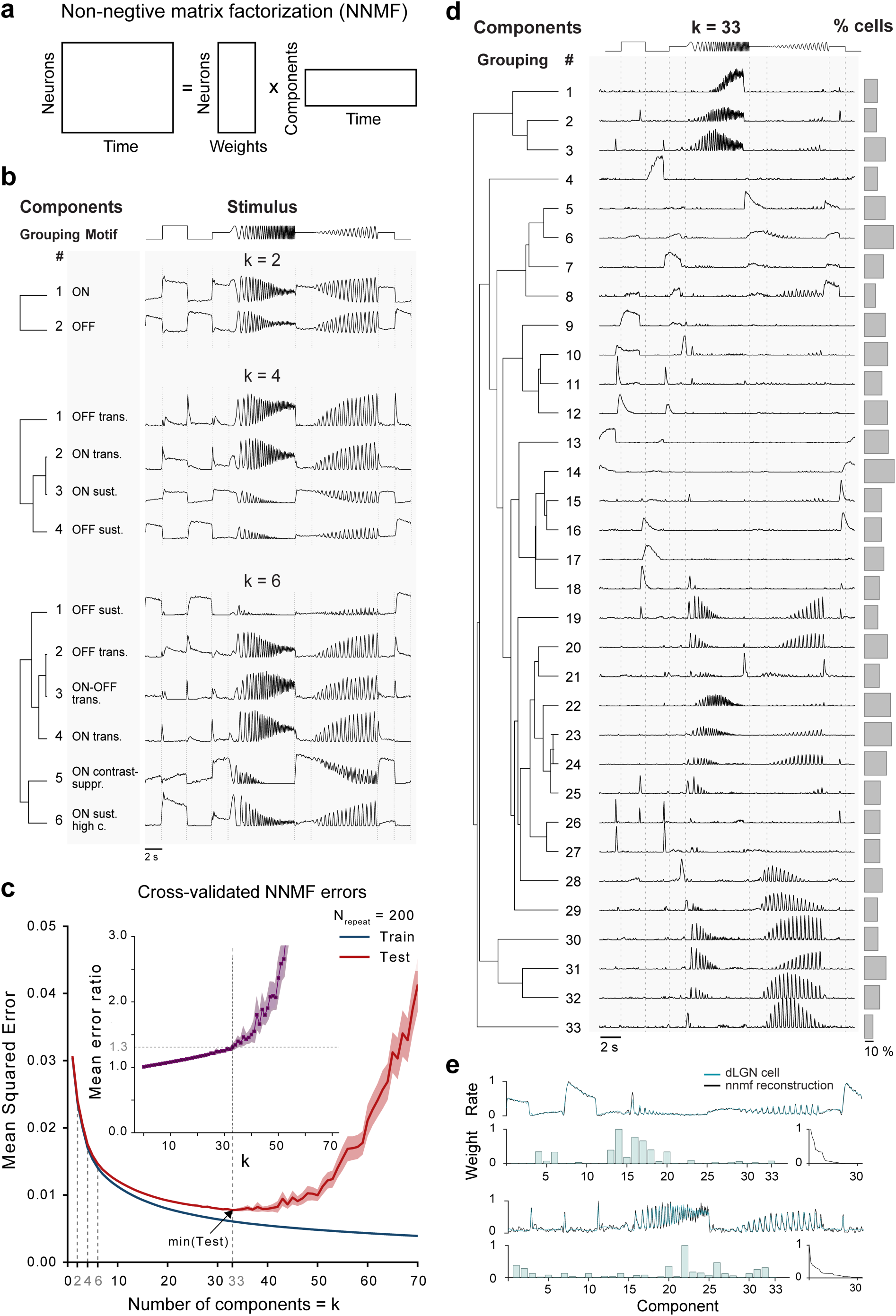
| Decomposition of the dLGN population response. **a,** Schematic of the non-negative matrix factorization (NNMF), used here to extract a set of time-varying visual response components, from which cell responses can be reconstructed as a weighted sum. **b,** Example NNMF models with k = 2, 4 and 6 components. Top: A model with two components clearly factorizes the data into an ON- and an OFF-component, which, at closer inspection, also differ in phase for the frequency and contrast modulation parts of the stimulus. Middle: Allowing four components introduces ON-transient and OFF-transient features in addition to sustained ON- and OFF-features. Bottom: With six components, components also reflect ON-OFF transient, as well as, ON-contrast suppressed features. **c,** Mean squared errors (MSE) as a function of rank (number of model components k) both for training (blue) and test (red) sets in NNMF model cross-validation (n_repeats_ = 200; lines, mean; shaded areas, CI_95_). NNMF cross-validation was used to determine the mathematically optimal number of components (k = 33) as the location of the minimum mean MSE_test_. Inset: MSE ratio (MSE_test_ / MSE_train_) per rank (line: mean; shaded area: CI_95_). **d,** dLGN response components as computed by NNMF (middle), organized according to a hierarchical cluster tree with an optimized leaf order (left), and percentage of neurons with weight of respective component > 0 (right). **e,** Top: Examples of dLGN cell responses (black) and their NNMF reconstruction (blue). Bottom: Weights used for reconstruction for each component (left), sorted in descending order (right).

Besides this rich representation of luminance steps, temporal frequencies and changes in contrast in the overall dLGN population, 82/443 (18.5%) dLGN neurons only displayed weakly modulated responses to the chirp stimulus, despite having robust and consistent responses to the drifting gratings presented before and afterwards (Supp. Fig. 4). This combination of response patterns is consistent with these dLGN neurons preferring local variations in luminance instead of full-field uniform patterns, as is the case for some RGC types ^3^. This suggests that the overall diversity of LGN responses may be larger than reported here. Together, this shows that the response diversity observed at the level of dLGN-p RGCs is also present in dLGN.

### Modelling dLGN responses as linear combinations of RGC inputs

Next, we combined the dLGN-p RGC dataset (Fig. 2) and the dLGN dataset (Fig. 3) to study how the dLGN responses are computed from the retinal output. We first accounted for the differences in recording methods by convolving the dLGN spiking responses with the OGB-1 Ca^2+^ indicator kernel (Supp. Fig. 2). We then used a linear model – constrained to have non-negative weights - to predict dLGN responses as a sum of weighted RGC inputs (Fig. 5a, Supp. Fig. 1). For prediction, we used the RGC-all cluster means ^3^ that were assigned at least two dLGN-p cells. Each dLGN recording consisted of multiple trials (stimulus repetitions, n = 10–30), which were divided into a training and a test set (50 % / 50 %). The model was evaluated using repeated random sub-sampling cross-validation with 1, 000 repetitions, where the weights were fitted on the training set in each repetition and prediction quality was assessed on the test set. The reported weights represent mean values across the repeats.

**Figure 5.**
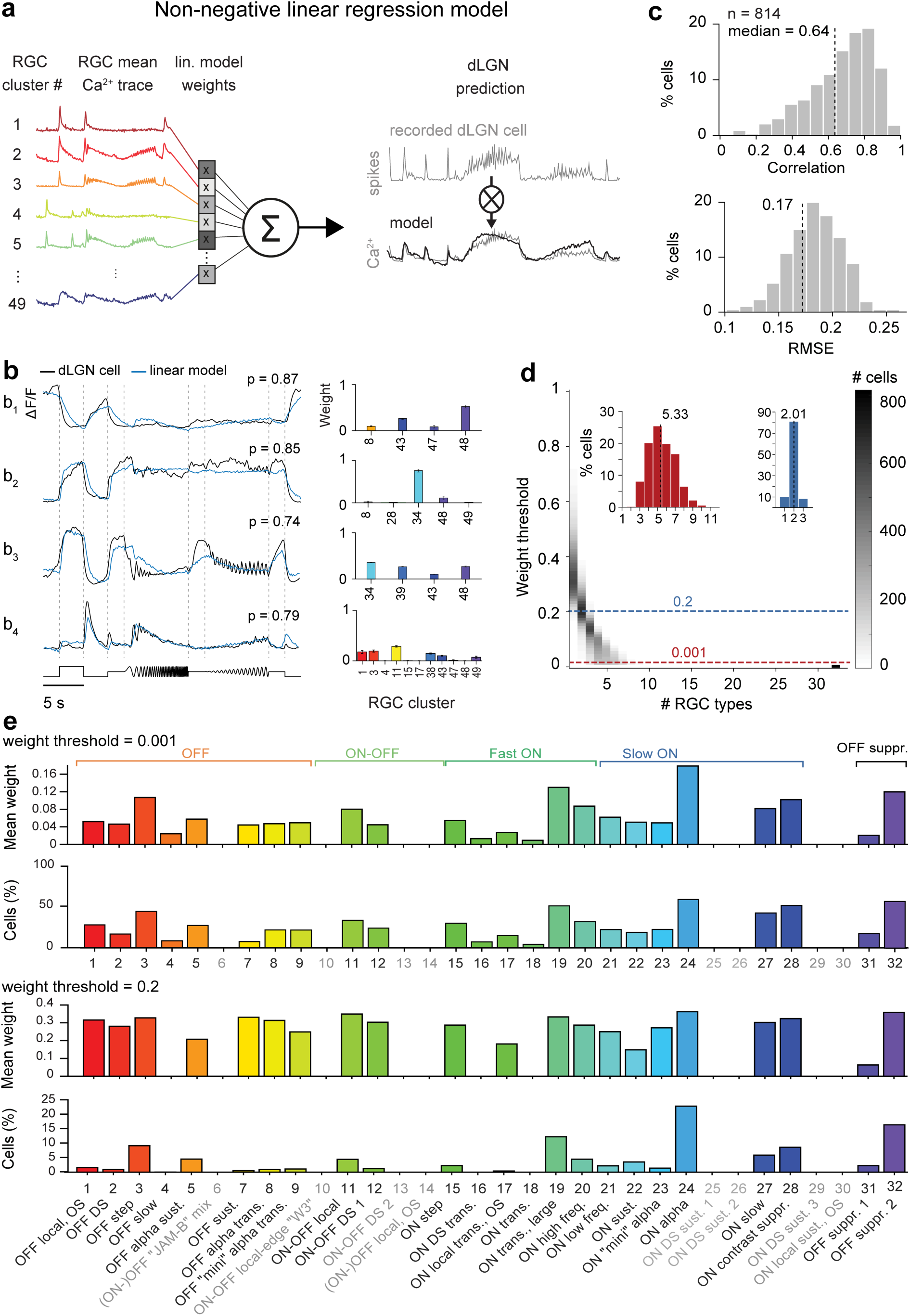
| Functional modelling of retinogeniculate convergence. **a**, Illustration of the linear model, which predicts responses of a dLGN neuron, convolved with a Ca^2+^ indicator kernel, as a linear combination of weighted dLGN-p RGC cluster inputs. **b**, Example responses of dLGN neurons (black) and predictions by the linear model (blue), along with the relative weights > 0.001 of RGC clusters used for modelling of dLGN responses. **c**, Prediction quality. Top: correlation between actual response and predicted response for all dLGN neurons (n = 814). Bottom: distribution of root mean squared errors **d**, Distribution of number of RGC groups used for reconstruction of dLGN neurons' responses, as a function of threshold for considering an input functionally relevant. **e**, Distribution of the percentage of dLGN neurons relying for functional reconstruction on the respective RGC groups from ^3^, along with the average weight (top: weight threshold = 0.001, bottom: weight threshold = 0.2). RGC types not projecting to dLGN are greyed-out.

This excitation-only linear feedforward model successfully reconstructed responses of most dLGN neurons, including transient ON, sustained ON and OFF, and suppressed cells (Fig. 5b). Typically, the responses of dLGN neurons could be predicted using only a few non-zero weights (weight > 0.001, Fig. 5b_1–3_); some cells were predicted by a combination of one dominant and few weaker RGC inputs (Fig. 5b_2_), and even others were best modelled with responses from many RGC clusters (Fig. 5b_4_). Across dLGN neurons, prediction quality was high (average correlation: 0.64, average RMSE: 0.17, Fig. 5c).

Using this model, we next asked how many RGC types provided input to mouse dLGN neurons and found that the answer directly depends on the threshold used for considering an input as functionally relevant (Fig. 5d). Choosing a low threshold, the responses of dLGN neurons (n = 814) could be predicted by the weighted input of on average 5 RGC groups (Fig. 5d, left inset, mean = 5.33, median = 5.19, range = 2–10, middle 90 % range = 3–8). Choosing a more conservative threshold for accepting functionally relevant inputs, in turn, identified on average 2 most dominant RGC groups (Fig. 5d, right inset, mean = 2.01). Together, these results indicate that the responses of many dLGN neurons, at least on the temporal scale of Ca^2+^ transients and for the presented set of visual stimuli, can be explained to a large extent by excitatory feedforward processing, with limited functional convergence.

Finally, we explored which of the previously identified dLGN-p RGC groups were used for prediction in the model (Fig. 5e). For the liberal weight threshold of 0.001, we found that the majority (24/32) of these groups were used by the model (Fig. 5e, top), with some of them contributing more frequently than others. Interestingly, some of the groups contributing most strongly belonged to those RGC groups that were significantly overrepresented among the dLGN-p RGCs (see also Fig. 2d), although information about the relative frequency of the projections was not used for inference. As expected, imposing a more conservative weight threshold resulted in an even more pronounced pattern of often-used RGC groups (Fig. 5e, bottom), while their relative weights tended to equalize. Together, our modelling approach suggests that these RGC types have a significant functional role in the processing of visual information along the retinogeniculate pathway.

## DISCUSSION

We functionally characterized the population of dLGN-projecting RGCs as well as LGN neurons and provide a quantitative account of the functional connectivity between RGC types and dLGN neurons. We present three main findings. First, combining a retrograde viral transduction approach and two-photon Ca^2+^ imaging of RGC responses, we show that the majority of previously functionally identified RGCs types projects to dLGN. Second, decomposing dLGN responses into their elementary response components revealed a rich diversity of geniculate visual representations, similar to that of RGCs. Third, we demonstrate that the responses of individual dLGN cells can be modelled as a linear combination of responses of on average 5 RGC types, amongst them 2 with the most dominant functional impact.

### Retrograde-labeling of dLGN-projecting RGCs

Retrograde viral tracing with HSV-cre combined with two-photon Ca^2+^ imaging of RGCs proved to be a suitable tool to determine functional properties of the dLGN-p RGCs. HSV is known to have a strictly synaptic uptake mechanism ^31,32^, which prevents infection of axons passing nearby and not synapsing within the dLGN. This is an issue with commonly used retrograde tracers, such as DiO / DiI, horseradish peroxidase, fluorophore-conjugated latex microspheres, or cholera toxin ^35^. In addition, our observation that GCaMP6f expression in the retina was limited to spatially confined regions is consistent with the interpretation that HSV infected neurons were restricted to the retinotopically corresponding region in dLGN instead of labeling passing RGC axons, which would likely have resulted in a larger spread of RGC locations.

### Functional classification of dLGN-projecting RGCs

Integrating the responses of dLGN-p RGCs with the functional classification obtained for the entire population of RGCs ^3^, revealed that around 75% of all functional RGC types innervate the dLGN. For example, ON alpha RGCs and OFF transient alpha RGCs ^39,40^ could be found more frequently among the dLGN-p cells than in the general RGC population. In agreement, ON sustained alpha RGCs were shown to target the core region of the dLGN ^41–43^ and also OFF transient alpha RGCs project to dLGN (for a complete comparison to the literature, see Supplementary Table 1). OFF suppressed RGCs, also referred to as Suppressed-by-Contrast (SbC-) RGCs, were also overrepresented in our dLGN-p dataset. Such cells typically exhibit a high baseline firing rate which is suppressed by light stimulation - a type of response that has been recorded along all stations of the mouse retino-geniculo-cortical pathway ^26,44,45^, suggesting a dedicated early visual pathway signaling uniformity of the visual field ^46–48^ and/or controlling contrast gain ^49^.

Certain RGC types like W3 RGCs or sustained ON DS RGCs are underrepresented among dLGN-p cells, in agreement with the literature on projections to dLGN ^50,51 52–54^. In two cases, our data seem add odds with the previous literature: None of the dLGN-p RGCs in our study were classified as JAM-B cell ^55^, which are known to project to dLGN. This is likely because JAM-B cells do not respond well to our set of visual stimuli; consistent with ref ^3^, where JAM-B cells were assigned to a “mixed” RGC group. Also, OFF sustained alpha RGCs were overrepresented among dLGN-p cells, although RGCs marked in the transgenic W7-line thought to correspond to OFF sustained alpha RGCs have been shown not to project to dLGN ^51^. Interestingly, there was a cluster in the OFF sustained group which was not assigned any dLGN-p cells, possibly indicating that this subgroup of cells corresponds to W7A. Alternatively, W7A RGCs are a different RGC type - in this case, all four alpha RGC types (ON-sustained, OFF-sustained, ON-transient, OFF-transient) ^40^ would project to dLGN in our data (the ON transient RGC likely corresponds to group 19), splitting the visual signal on the way to dLGN into four channels arranged symmetrically with respect to polarity and kinetics ^39^–^40^–^56^.

### Components of the dLGN population response

Consistent with its rich retinal input, we found that the dLGN population responses can be factored into more than 30 components. The specific number certainly depends on exactly which method is used for determining the optimal rank of the model. We tested several of these (AIC, BIC, randomization, cross-validation), and found none of them to suggest numbers as low as the classical notion of 3 parallel pathways (reviewed in ^57^). Instead, the high number of components points towards a larger diversity of visual features encoded by dLGN neurons than commonly appreciated, at least in mice. This interpretation is supported by recent studies reporting “non-classical” responses in rodent dLGN ^25–27,58–60^, rabbit dLGN ^61^ and the koniocellular layers of primate dLGN ^62–64^, such as direction and orientation selectivity, and binocularity. While our factorization approach reveals an elaborate visual representation at the level of the dLGN, it is still an open question how many distinct types of dLGN neurons exist. A previous clustering study of dLGN responses to a large stimulus set estimated the number of functional clusters to be six ^26^. While the identified clusters undoubtedly have correlated response profiles, it is unclear to which degree their borders are strictly demarcated or instead the clusters rather form a continuum. We also attempted to cluster our dLGN data, similar to ^26^, but - in light of the large diversity we observed in our recordings - we were not convinced by the separation of the response clusters based on the limited number of recordings.

### Recombination of RGC channels in dLGN

In previous work, the question of retinogeniculate connectivity has been addressed from two different angles. Most studies focused on the absolute number and strength of RGC inputs to single dLGN neurons, whereas fewer studies also assessed convergence of retinogeniculate inputs in terms of RGC types.

Estimates of the number of individual RGCs providing input to a single dLGN relay cell differ widely between studies (between one and ninety), with the discrepancy likely arising from differences in model species, age, and method of investigation. Both *in-vitro* and *in-vivo* physiology studies concluded that dLGN neurons typically receive input from 1–3 dominant RGCs, with the possibility of additional, weaker inputs in cats and rodents ^8,9,15,65–70^. For example, in adult mice, a recent paper reported three dominantly driving inputs, with an average total number of 10 RGC inputs per dLGN relay cell. This estimate was derived from the so-called “fiber fraction technique”, where the size of the EPSP evoked by stimulation of a single fiber in the optic tract is compared to the EPSP evoked by stimulating all fibers. Recent studies using EM reconstructions ^71,72^ and mono-transsynaptic rabies virus tracing ^17^ in mice suggested the possibility that a rather large number (up to 91) of RGCs can converge onto single dLGN neurons. These numbers appear high compared to previous estimates, possibly because many of the synapses identified structurally have a low weight ^73^. Furthermore, discrepancies between estimates may be partially due to differences in the age of the mice, where functional refinement including synapse elimination and axon retraction occurs between the ages of P9 and P60 ^65,68,74^.

Similarly, the estimated number of RGC types projecting to a single dLGN relay cell varies across studies, and will likely differ depending on model species (reviewed in ref ^72^). In cats, where dLGN is organized into functionally distinct layers, paired recordings of RGCs ^8,75^ or S-potentials ^9^ and dLGN relay cells have generally found an increase in connection probability with increasing similarity of RF properties. In particular, monosynaptic connections between RGCs and dLGN relay cells of opposite polarity have been rarely observed^8,9,14,15^, indicating that, at least in cats, convergence of RGC types might be low. The most direct estimates of convergence amongst RGC types to date has been obtained in mice, where single dLGN relay cell initiated mono transsynaptic rabies virus tracing combined with morphological analysis of labeled RGCs revealed that some dLGN relay cells received input from RGCs of one or mostly one type, while others received input from up to nine different cell types ^17^. Our estimates based on our computational modelling agree with both findings and highlight the importance of considering which inputs are functionally relevant. Our results with a very liberal weight threshold indicate that response prototypes from 3–8 RGC types are required to account for single cell dLGN responses, compatible with the recent anatomical estimates based on EM reconstructions and mono-transsynaptic rabies virus tracing ^17,19^. Raising the threshold to weights more likely having functionally relevant impact, reduces the estimated number of RGC types providing inputs to mouse dLGN neurons to two, consistent with the large body of literature on functional convergence in the retino-geniculate pathway.

Considering its simplicity, our feedforward model of dLGN responses performed well. This is surprising, because the model does not consider known components of geniculate circuits, such as local inhibitory interneurons ^76^, cortico-geniculate feedback ^77^, and neuromodulatory influences ^78,79^. Potential explanations for the success of the model despite disregarding these essential circuits include the spatial simplicity of our stimulus set and the low-pass nature of our signals. Indeed, local inhibition is known to increase push-pull mechanisms in dLGN, thereby enhancing temporal and spatial contrast ^76,80^. These mechanisms will likely play a larger role in the processing of localized or more complex stimuli, such as natural scenes ^81^, than in the full-field stimulus used here. Similarly, performing the model predictions with the temporal resolution of Ca^2+^ signals likely smoothens over differences in firing patterns associated with behavioral state and neuromodulation ^44,82,83^. In the future, it will be essential to explore model predictions with more complex visual stimulation and signals of higher temporal resolution.

## METHODS

All procedures complied with the European Communities Council Directive 2010/63/EC and the German Law for Protection of Animals, and were approved by local authorities, following appropriate ethics review.

### Functional characterization of dLGN-projecting RGCs

#### Animals and virus injection

For all experiments, we used 8 to 12 week-old animals (either sex) of the Ai95D reporter line (B6; 129S- Gt(ROSA)26Sor^tm95.1(CAG-GCaMP6f)Hze^/J; JAX 024105). Ai95D features a floxed-STOP cassette preventing transcription of the genetically-encoded Ca^2+^ indicator GCaMP6f ^29^. Stereotactic injection of a Cre-encoding Herpes-Simplex-Virus 1 (hEFIα-cre, MIT Vector Core, Cambridge, USA) into the dLGN resulted in retrograde Cre-recombinase expression in dLGN-projecting (dLGN-p) RGCs, where Cre-recombinase, in turn, removed the LoxP sites and activated GCaMP6f expression ^84^.

The surgical procedures have been described previously ^82,85^. In brief: Mice were fixed in a stereotactic frame (Neurostar, Tübingen, DE) and anaesthetized using an isoflurane–oxygen mixture (4.0% induction, 1.2% maintenance) throughout the entire surgery. At the beginning of the surgical procedure, atropine (Atropine sulphate, 0.3 mg/kg, sc, Braun, Melsungen, DE) and analgesics (Buprenorphine, 0.1 mg/kg, sc, Bayer, Leverkusen, DE) were administered, and the eyes were protected continuously with an eye ointment (Bepanthen, Bayer, Leverkusen, DE). The animal's temperature was kept constant at 37°C via a closed loop temperature control system for small rodents (HD, Hugo Sachs Elektronik-Harvard Apparatus GmbH, March-Hugstetten, DE).

After a midline scalp incision, a small hole was made with a dental drill (Sinco, Jengen, DE) over the LGN located in the left hemisphere, 2.5 mm posterior to the bregma and 2.3 mm lateral from the midline. The virus was loaded in a sharp micropipette (GB150F-8P, Science Products, inner tip diameter 20–25 μm, Hofheim, DE) connected through a 10 μl Hamilton syringe (Hamilton Robotics, Reno, USA) to an Aladdin syringe pump (AL-1000, WPI Germany, Berlin, DE). A volume of 20–40 nl of virus was injected at a depth of 2.7 mm. The pipette was left in place for an additional 5 min to allow for viral diffusion. Antibiotics (Baytril, 5 mg/kg, sc, Bayer, Leverkusen, DE) and a longer lasting analgesic (Carprofen, 5 mg/kg, sc, Rimadyl, Zoetis, Berlin, DE) were administered continuously for 3 days post-surgery. Two-photon Ca^2+^ imaging was carried out 3 weeks after viral injection.

#### Perfusion and retinal tissue preparation

Animals were housed under a standard 12 h day/night rhythm. Before perfusion and two-photon imaging, animals were dark-adapted for ≥1 h, and then deeply anaesthetized with a lethal dose of sodium pentobarbital (Narcoren, 400 mg/kg, injected intraperitoneally, Böhringer Ingelheim, Ingelheim, DE). When the animal reached the asphyxia stage and complete paralysis, the eyes were enucleated, and the mouse was transcardially perfused with 0.2 M sodium phosphate buffered saline (PBS), followed by 4% paraformaldehyde (PFA) solution in PBS. The brains were postfixed in PFA for 24 hours at 4° and then stored in PBS.

The eyes were dissected in carboxygenated (95% O_2_, 5% CO_2_) extracellular solution containing (in mM): 125 NaCl, 2.5 KCl, 2 CaCl_2_, 1 MgCl_2_, 1.25 NaH_2_PO_4_, 26 NaHCO_3_, 20 glucose, and 0.5 L-glutamine (pH 7.4). The retina was extracted from the eyecup and flat-mounted onto a ceramic filter (Anodisc #13, 0.2 μm pore size, GE Healthcare, Buckinghamshire, UK) with the ganglion cell layer (GCL) facing up and transferred to the recording chamber of the microscope, where it was continuously perfused with carboxygenated solution at ~36 °C. In all experiments, ~0.1 μM Sulforhodamine-101 (SR101, Sigma, Steinheim, DE) was added to the extracellular solution to reveal blood vessels and any damaged cells in the red fluorescence channel of the microscope ^86^. All procedures were carried out under dim red (> 650 nm) illumination.

#### Histological reconstruction of injection sites

To verify the injection site within the dLGN, we used histological reconstructions. Brains were sliced for coronal sections (50 μm) using a vibratome (Microm HM 650 V, Thermo Fisher Scientific, Waltham, Massachusetts, USA) and mounted on glass slides with DAPI-containing mounting medium (Vectashield DAPI, Vector Laboratories Ltd, Peterborough, UK), which labels the cell nuclei, and cover-slipped. Brain slices were inspected using a Zeiss Imager.Z1m epi-fluorescent microscope (Zeiss, Oberkochen, DE) for the expression of the retrogradely transported HSV-1, visualized by the expression of the GCaMP6f protein containing the enhanced green fluorescent protein (eGFP).

#### Two-photon Ca^2+^ imaging and light stimulation

We used a MOM-type two-photon microscope (designed by W. Denk, MPI, Martinsried; purchased from Sutter Instruments/Science Products, Hofheim, Germany). Design and procedures were described previously ^3,86^. In brief, the system was equipped with a mode-locked Ti:Sapphire laser (MaiTai-HP DeepSee, Newport Spectra-Physics, Darmstadt, Germany) tuned to 927 nm, two fluorescence detection channels for GCaMP6f (HQ 510/84, AHF/Chroma Tübingen, Germany) and SR101 (HQ 630/60, AHF), and a water immersion objective (W Plan-Apochromat 20x/1.0 DIC M27, Zeiss, Oberkochen, Germany). For image acquisition, we used custom-made software (ScanM, by M. Müller, MPI, Martinsried, and T. Euler) running under IGOR Pro 6.37 for Windows (Wavemetrics, Lake Oswego, OR, USA), taking 64 × 64 pixel image sequences (7.8 frames/s) for activity scans or 512 × 512 pixel images for high-resolution morphology scans.

For light stimulation, we focused a DLP projector (K11, Acer) through the objective, fitted with band-pass-filtered light-emitting diodes (LEDs) (“green”, 578 BP 10; and “blue”, HC 405 BP 10, AHF/Chroma) to match the spectral sensitivity of mouse M- and S-opsins. LEDs were synchronized with the microscope's scan retrace. Stimulator intensity (as photoisomerization rate, 10^3^ P*/s/cone) was calibrated as described previously ^86^ to range from 0.6 and 0.7 (black image) to 18.8 and 20.3 for M- and S-opsins, respectively. An additional, steady illumination component of ~10^4^ P*/s/cone was present during the recordings because of two-photon excitation of photopigments (for detailed discussion, see ^86,87^). For all experiments, the tissue was kept at a constant mean stimulator intensity level for at least 15 s after the laser scanning started and before light stimuli were presented.

Four types of light stimuli were used (Fig. 1d, top, ref ^3^): (*i*) Full-field (800 × 600 μm) “chirp” stimuli consisting of a bright step and two sinusoidal intensity modulations, one with increasing frequency (0.5–8 Hz) and one with increasing contrast; (*ii*) 0.3 × 1 mm bright bar moving at 1 mm s^-1^ in eight directions; (*iii*) alternating blue and green 3-s flashes; and (*iv*) binary dense noise (20 × 15 matrix with 40 μm pixel-side length; each pixel displayed an independent, balanced random sequence at 5 Hz for 5 minutes) for space-time receptive field mapping. All stimuli, except (*iii*), were achromatic, with matched photo-isomerization rates for mouse M- and S-opsins.

#### Data analysis (dLGN-projecting RGCs)

The data analysis was performed using IGOR Pro (Wavemetrics, Lake Oswego, OR, USA), MATLAB (The Mathworks, Natick, Massachusetts, USA) and Python (distribution by Anaconda Inc., Austin, TX) using methods described previously ^3^.

#### Pre-processing

Regions of interest (ROIs) were manually drawn around the GCaMP6f-expressing somata in the recording fields. The Ca^2+^ traces for each ROI were extracted (as ∆F/F) using the image analysis toolbox SARFIA for IGOR Pro ^88^. A stimulus time marker embedded in the recorded data served to align the Ca^2+^ traces relative to the visual stimulus with a temporal precision of 2 ms. For this, Ca^2+^ traces were up-sampled to 512 Hz, and the timing for each ROI was corrected for sub-frame time-offsets related to the scanning. The Ca^2+^ traces were then de-trended using a high-pass filtering above ~0.1 Hz and resampled to 7.8 Hz. For all stimuli except the dense noise (for RF mapping), the baseline was subtracted (median of first eight samples), median activity *r(t)* across stimulus repetitions computed (typically three to five repetitions) and normalized such that *max_t_*(|*r(t)*|) = 1.

#### Direction and orientation selectivity

To extract time course and directional tuning of the Ca^2+^ response to the moving bar stimulus, we performed a singular value decomposition (SVD) on the *T* by *D* normalized mean response matrix *M* (times samples by number of directions; *T* = 32; *D* = 8):

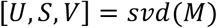

This procedure decomposes the response into a temporal component in the first column of *U* and a direction dependent component or tuning curve in the first column of *V*, such that the response matrix can be approximated as an outer product of the two:

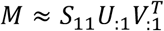

An advantage of this procedure is that it does not require manual selection of time bins for computing direction tuning but extracts the direction-tuning curve given the varying temporal dynamics of different neurons.

To measure direction selectivity (DS) and its significance, we projected the tuning curve *V*_:1_ on a complex exponential *ϕ_k_*= exp(*iα_k_*), where *α_k_* is the direction of the *k_th_* condition:

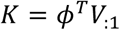

This is mathematically equivalent to computing the vector sum in the 2D plane or computing the power in the first Fourier component. We computed a DS index as the resulting vector length:

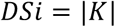

correcting for the direction spacing. We additionally assessed the statistical significance of direction tuning using a permutation test ^89^. To this end, we created surrogate trials (that is, stimulus repetitions) by shuffling the trial labels (that is, destroying any relationship between condition and response), computed the tuning curve for each surrogate trial and projected it on the complex exponential *ϕ*.

Carrying out the procedure 1,000 times generated a null distribution for K, assuming no direction tuning. We used the percentile of the true K as the P value for direction tuning ^3^.

Orientation selectivity (OS) was assessed in an analogous way. However, we used the complex exponential *ϕ_k_*= exp(*2iα_k_)*, corresponding to the second Fourier component.

#### Signal deconvolution

Comparing neural activity measured with different methods (i.e. spikes vs. Ca^2+^ signals; different Ca^2+^ indicators) is a non-trivial task. To assign dLGN-p RGCs to previously characterized RGC types, we needed account for the fact that the different Ca^2+^ indicators used (OGB-1 vs. GCaMP6f) have different kinetics ^29^. We decided to convert both signals to a “common currency”, be deconvolving both signal types using Ca^2+^ kernels calculated for each indicator separately using Ca^2+^ recordings of multiple ROIs (n_OGB-i_ = 327; n_GCaMP6f_ = 19) to the white noise stimulus, and averaging thresholded Ca^2+^ peak events (>80% of the maximum normalized activity). The kernel area under the curve was normalized to 1. (Supp. Fig. 2).

#### Assigning dLGN-projecting RGCs to previously characterized RGC types

The pre-processed ROI traces of dLGN-p RGCs (n = 251) were assigned to the functional RGC population clusters reported by ^3^, by identifying for each dLGN-p cell the cluster with the best matching response properties. After deconvolution with the respective Ca^2+^ kernel (see above), we calculated the linear correlation coefficients between a dLGN-p cell's mean trace (over trials) and all cluster mean traces (over all cells in an RGC population cluster) for the chirp and the moving bar stimuli. To combine the information about stimulus-specific correlations and stimulus-specific cell response quality, we generated an overall match index (Mi) of each dLGN-p cell to all RGC population clusters:

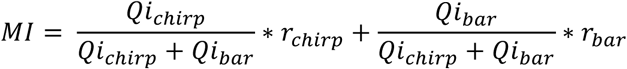

Finally, each dLGN-p cell was assigned to the cluster with the highest Mi.

#### Comparing relative cluster proportions in dLGN-p RGCs and RGC-all

We used a binomial test to assess the statistical significance of deviations of cluster cell numbers in the dLGN-p RGCs subpopulation compared to the RGC-all population. Starting with a total dLGN-p RGC cell number *N_dLGN-p RGC_* = 251, for each dLGN-p RGC cluster *i*, we computed the test parameters *k_i_* = “number of cells cluster *i* in dLGN-p RGC population”, and *p_i_* = “proportion of cells in cluster *i* in the RGC-all population” *(N_RGC-all_* = 5024), where p_i_ would be the expected proportion, or probability, of cells in that cluster if it were the same as in the RGC-all cluster. We then performed a two-tailed binomial test of obtaining a cell number deviation that extreme given the null hypothesis of equal cluster cell numbers in the two populations. After p-value correction for false discovery rate (FDR, Benjamini-Hochberg), p-values were considered significant at the significance level alpha = 0.01.

For the log_2_-ratio (% dLGN-p RGCs / % RGC population) plot, we used additive smoothing of the histogram, i.e. by adding one to each group's cell count n per default and normalizing the group cell counts by n+k, where k is the number of clusters.

#### Linear feedforward model

For modelling dLGN responses as a linear combination of weighted RGC types, we used RGC-all cluster means ^3^ only of those clusters that were assigned at least two cells from our dLGN-p RGC data. (Supp. Fig. 1). The spike dataset recorded in dLGN was down-sampled and convolved with an artificial OGB-1 kernel to allow for a direct comparison of the dLGN traces with the dLGN-p RGC Ca^2+^ responses. We then modelled dLGN responses as a linear combination of weighted RGC inputs (Supp. Fig. 1). The weights were computed, using a linear-regression algorithm *(lsqlin*, MATLAB) with a non-negativity constraint, and then applied to their respective RGC population cluster responses, yielding the optimal prediction of the dLGN cell response. The non-negativity constraint is motivated by the non-negativity of feedforward excitatory connections between RGCs and LGN. The model performance was cross-validated using repeated random sub-sampling with 1,000 repetitions. The trials were randomly shuffled and divided into a training set (50% trials) and a validation set (50% trials). Both sets thus contained unique trials with no duplicates. The training set was used to compute the weights and the performance of the model was evaluated on the validation set. In the end, a particular dLGN cell was modelled with the mean weights across all repeats, and the performance of the model represents the mean value across the repeats.

### Functional characterization of dLGN responses

#### Animals and surgical procedures

For all experiments, we used 8- to 12-week-old wild type mice (C57BL/6J) of either sex. For the initial surgery procedures refer to the section “Animals and virus injection”.

After a midline scalp incision and skin removal, a drop of H_2_O_2_ (3 %) was applied on the surface of the skull for the removal of tissue residues. A custom lightweight aluminum head post was placed on the posterior skull using OptiBond FL primer and adhesive (Kerr dental, Rastatt, DE) and Tetric EvoFlow dental cement (Ivoclar Vivadent, Ellwangen, DE). Miniature ground and reference screws (00–96 × 1/16 stainless steel screws, Bilaney) soldered to custom-made connector pins were placed bilaterally over the cerebellum. A well of dental cement was formed to hold the silicone elastomer sealant Kwik-Cast (WPI Germany, Berlin, DE) covering the skull. Antibiotics (Baytril, 5 mg/kg, sc, Bayer, Leverkusen, DE) and a longer lasting analgesic (Carprofen, 5 mg/kg, sc, Rimadyl, Zoetis, Berlin, DE) continued to be administered for 3 days post-surgery.

After recovery, animals were familiarized with a simulation of the experimental procedures in multiple training sessions until they were deemed comfortable with the conditions. Before experiments, a craniotomy (ca. 1 mm^2^) was performed over dLGN (2.3 mm lateral to the midline and 2.5 mm posterior to bregma), which was re-sealed with Kwik-Cast (WPI Germany, Berlin, DE). Experiments started one day after craniotomy and were continued on consecutive days as long as electrophysiological signals remained of high quality.

#### In-vivo multisite extracellular recordings

Our experimental configuration for in-vivo recordings was based on ^90^. The mouse was head-fixed and could run freely on an air-suspended styrofoam ball while stimuli were presented on a gamma-corrected LCD screen (Samsung SyncMaster 2233). Extracellular neural signals were recorded 2.5 mm posterior from bregma and 2.3 mm lateral from midline through a small craniotomy window over dLGN with 32-channel edge silicon probes (Neuronexus, A1×32Edge-5mm-20–177-A32, Ann Arbor, USA). Neurons were verified as belonging to the dLGN based on the characteristic RF progression from top to bottom along the electrode shank (Fig. 3c), the preference for high temporal frequencies, and a high prevalence of F1 responses to drifting gratings ^24,26^. Ball movements were registered at 90 Hz by two optical mice connected to an Arduino-type microcontroller (http://arduino.cc). Eye movements were monitored under infrared light illumination (Guppy AVT camera, frame rate 50 Hz, Allied Vision, Exton, USA).

#### Visual stimulation

We used custom software (EXPO, https://sites.google.com/a/nyu.edu/expo/home) to present visual stimuli on a gamma-calibrated liquid crystal display (LCD) monitor (Samsung SyncMaster 2233RZ; mean luminance 50 cd/m^2^, 60 Hz) at 25 cm distance to the animal's right eye. Four types of light stimuli were presented: (*i*) a "contrast stimulus” to measure the contrast response function, consisting of drifting sinusoidal gratings at a single orientation and 12 different randomly interleaved contrasts for 2 secs with 5 sec pauses between trials. (*ii*) the full-field chirp stimulus (see RGC section). (*iii*) a spatial-temporal-frequency-orientation (STFO) stimulus to capture preferred tuning properties of a large number of neurons simultaneously; the stimulus consisted of drifting sinusoidal gratings with 8 orientations, 6 temporal (0.5, 1, 2, 4, 8, 16 cycles/sec) and 2 spatial frequencies (0.5, 0.15 cycles/°) (443 of 814 cells, Supp. Fig. 4). Trials were randomly interleaved and presented for 1 sec with 0.1 sec pauses between trials. The stimulus was shown at 100% contrast with the background at mean luminance. (*iv*) a sparse noise stimulus for receptive field mapping, consisting of white or black squares (5° visual angle) presented on a background of mean luminance (50 cd/m^2^). Squares were flashed for 200 ms each at every position on a 12 × 12 square grid of 60 degrees.

All light stimuli were presented in a full-field mode. In experiments measuring tuning curves, a blank screen condition (mean luminance) was included in all stimuli to estimate the spontaneous firing rate.

#### Data analysis (dLGN responses)

Data analysis was performed using Matlab (The Mathworks, Natick, Massachusetts, USA). Data were organized in a custom written schema using the relational database framework "DataJoint" ^91^ (https://github.com/datajoint/datajoint-matlab).

#### Unit extraction and spike sorting

Wideband extracellular signals were digitized at 30 kHz (Blackrock microsystems, Blackrock Microsystems Europe GmbH, Hannover DE) and analyzed using the NDManager software suite ^92^. The LFP was computed after downsampling to 1250 Hz. To isolate single neurons from the linear arrays, we grouped neighboring channels into 5 equally sized “virtual octrodes” (8 channels per group with 2 channel overlap for 32 channel probes). Using an automatic spike detection threshold ^93^ multiplied by a factor of 1.5, spikes were extracted from the high-pass filtered continuous signal for each group separately. The first three principal components of each channel were used for semi-automatic isolation of single neurons with KlustaKwik ^94^. Clusters were manually refined with Klusters ^92^. We assigned each unit to the contact with the largest waveform. Units were given a subjective quality score by the manual sorter, the firing rate, the cleanness of the refractory period, and the stability over time. To avoid duplication of neurons extracted from linear probe recordings, we computed cross-correlation histograms (CCHs, 1 ms bins) between neuron pairs from neighboring groups. Pairs for which the CCH's zero-bin was three times larger than the mean of non-zero-bins were considered to be in conflict. For each conflicting pair, the cell with the best score was kept. Conflicts across pairs were resolved by collecting all possible sets of cells and by keeping the set with the best total score.

#### Receptive field mapping

Receptive fields were mapped by reverse correlating unit activity to the sparse noise stimulus and fitting the center of a two-dimensional ellipse / 2D-Gaussian for both ON- and OFF-fields ^95^:

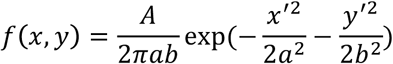

where *A* is the maximum amplitude, *a* and *b* are half-axes of the ellipse, and *x’* and *y’* are transformations of the stimulus coordinates *x* and y, considering the angle *θ* and the coordinates of the center (*xc*, *yc*) of the ellipse. For each contact, we computed a single RF center by averaging coordinates of the best-fit ON and OFF subfield (explained variance > 70%).

#### Contrast response function

Contrast responses were fitted with a hyperbolic ratio function ^96^:

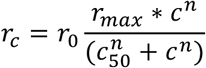

with baseline response *r*_0_, responsiveness *r_max_*, semi-saturation contrast *c*_50_, and exponent *n*.

#### Tuning

Orientation tuning curves were fitted with a sum of two Gaussians with peaks *θ_pref_* and *θ_pref_ − π* of different amplitudes *A*_1_ and *A*_2_ but equal width *σ*, with a constant baseline *r*_0_^97^. For each neuron, spatial and temporal frequency tuning curves were taken at its preferred direction; orientation and direction-tuning curves were taken at its optimal spatial and temporal frequencies.

***Direction selectivity*.** Direction selectivity index (DSI) was calculated as the ratio of

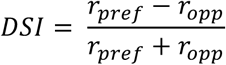

where *r_pref_* was the response at the preferred direction and *r_opp_* was the response at the opposite direction. We additionally assessed the statistical significance of direction tuning using a permutation test ^89^ as described above.

***Orientation selectivity*.** Orientation selectivity index (OSI) was computed as:

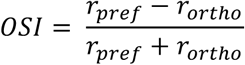

where *r_pref_* is the response to the preferred orientation and *r_ortho_* is the response to the orthogonal orientation^98^.

#### Other response measures

***Response quality index*.** To measure how well a cell responded to a stimulus (local and full-field chirp, flashes), we computed the signal-to-noise ratio

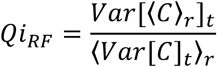

where *C* is the *T* by *R* response matrix (time samples by stimulus repetitions), while *<>_x_* and *Var[]_x_* denote the mean and variance across the indicated dimension, respectively ^3^. For further analysis, we used only cells that responded well to the chirp and/or to the moving bar stimulus *(Qi_chirp_ >* 0.45 or *Qi_DS_ > 0.6*, Supp. Fig. 1, cf. ^3^). Of the original n = 581 ROIs, n = 251 ROIs passed this criterion.

***Chirp quality index*.** To determine whether a neuron was visually driven by the full-field chirp, we separated the stimulus into two segments (e.g., separated a 32-sec stimulus into two 16 sec segments), and computed the average between-trial correlations (CCs) (responses binned at the stimulus frame rate) within segment and between segments. Only those cells that had significantly higher within-segment CCs (Qi_chirp_, P < 0.01, Wilcoxon rank sum test) and firing rate > 1 spike/s were considered to be visually responsive.

***STFO correlation value*.** To assure neuronal response stability and to determine, whether there were neurons that did not respond to the full-field chirp stimulus, we played the STFO stimulus directly before and after the chirp stimulus and performed linear regression analysis (Supp. Fig. 4). To determine how well the model predicts the data, we computed a correlation value *R:*

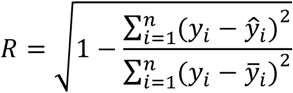

where ŷ represents the calculated values of *y* and 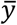 is the mean of y.

For further analysis, we used only cells that responded well to the chirp stimulus *(Qi_RF_ >* 0.05 and *Qi_chirp_* ≤ 0.001) or had a high STFO correlation value *(R* > 0.65, Supp. Fig. *1)*.

#### Non-negative matrix factorization (NNMF)

NNMF decomposes a matrix *A* into two low-rank non-negative matrices *U* and *V^T^*, representing elementary components and their weights ^37^. In the case of our dLGN cell responses, this translates into finding a set of time-varying visual response components from which cell responses can be reconstructed as a weighted combination of those components. Given a positive matrix *A* of size *N × M* (neuron x time) and a desired number of features *(K)*, the NNMF algorithm iteratively computes an approximation *A ~ UV^T^*, where *U* and *V^T^* are non-negative matrices with respective sizes *N × K* (neuron x weight) and *K × M* (component x time).

To determine the optimal number of components *K*, we applied NNMF cross-validation ^38^, as described by Alex H. Williams on http://alexhwilliams.info/itsneuronalblog/2018/02/26/crossval/ and implemented on https://gist.github.com/ahwillia/65d8f87fcd4bded3676d67b55c1a3954. Analogous to cross-validation in supervised learning settings, NNMF cross-validation splits the data into a training and a test set in order to compare model performance on both sets, and uses decreasing test set performance as a function of *K* as an indicator of model overfitting. In contrast to supervised settings, data entries cannot simply be held out for entire rows or columns as this would not allow to fit all parameters. Here, this problem was overcome by randomly holding out individual data entries from the data matrix and using the remaining data entries as training set^99^. Formally, we define a binary matrix *M*, which acts as a mask over our data. The results in the following optimization problem:

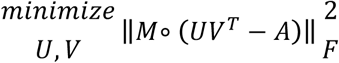

The optimization problem amounts to the low-rank matrix completion problem which was solved by alternating minimization of the above equation for *U* and *V^T^*. The model error can then be evaluated on the held-out data points as follows:

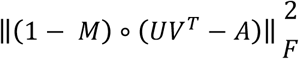

We fitted NNMF models on the training data for ranks (= number of components k) 1 to 70, and compared the mean squared error (MSE) for the training vs. the validation set. Due to the random model initialization for each run, this was done with n_repeats_ = 200. The optimal number of components was determined as the rank with the minimal average MSE over all runs (here, k = 33).

The resulting components were organized into a hierarchical cluster tree by a single linkage algorithm using Euclidean distances and the Ward's minimum variance method ^100^. The results were plotted using the dendrogram function and the leaf order was optimized using the Matlab function *optimalleaforder*.

### Histological reconstruction of recording sites

To verify recording sites from dLGN, we used histological reconstructions. Before recording from the dLGN, electrodes were coated with a red-shifted fluorescent lipophilic tracer (DiD; Thermo Fisher Scientific, Waltham, Massachusetts, USA). After the last recording session, mice were transcardially perfused and the brain fixed in a 4% paraformaldehyde phosphate buffered saline (PBS) solution for 24 hours and then stored in PBS. Brains were sliced for coronal sections (50 μm) using a vibratome (Microm HM 650 V, Thermo Fisher Scientific, Waltham, Massachusetts, USA) and mounted on glass slides with Vectashield DAPI (Vectashield DAPI, Vector Laboratories Ltd, Peterborough, UK), and coverslipped. Slices were inspected for DAPI and DiD presence using a Zeiss Imager.Z1m fluorescent microscope (Zeiss, Oberkochen, DE).

## SUPPLEMENTARY FIGURES

### 1. Overview of analysis steps

**Supplementary Figure 1.**
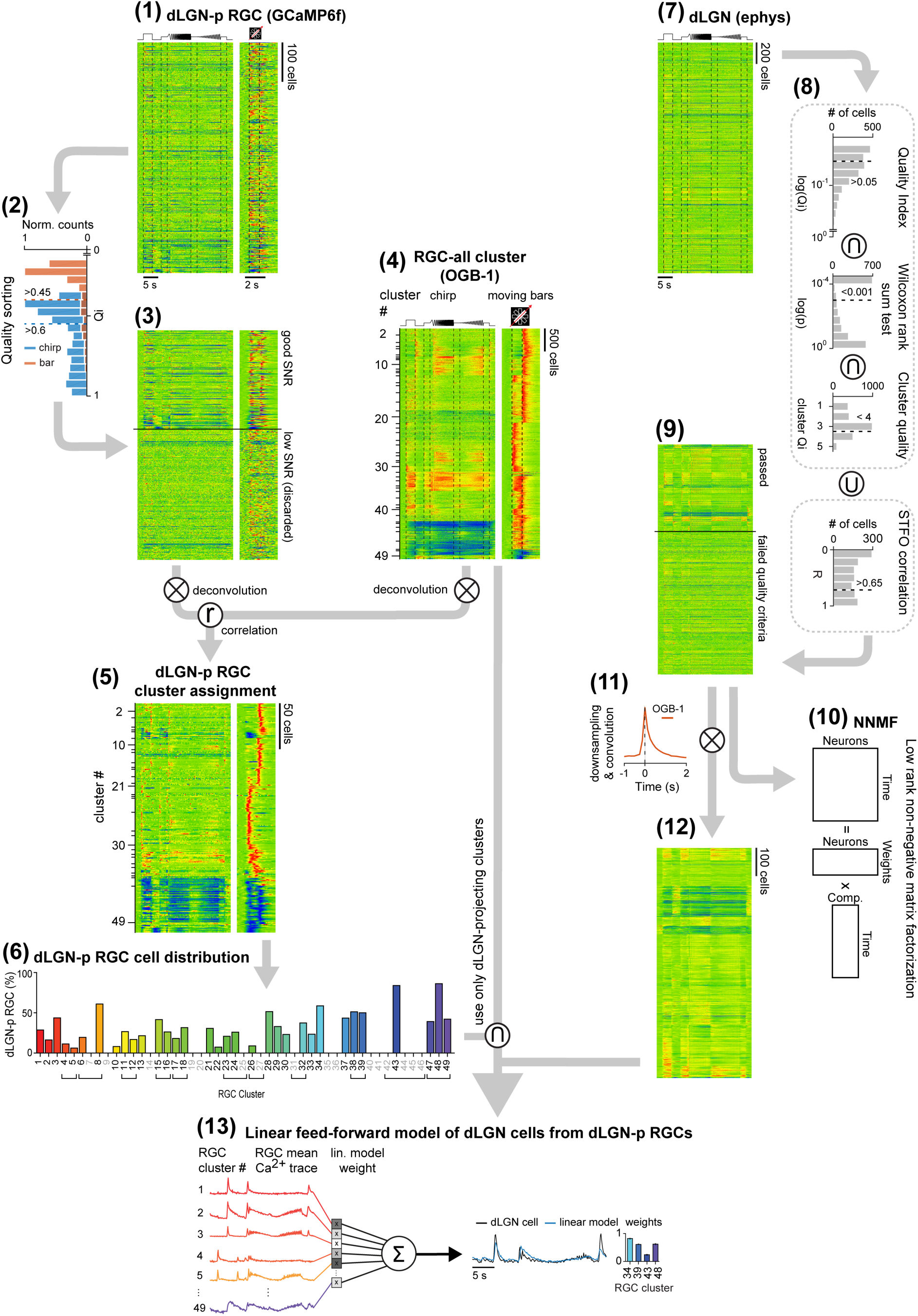
| Processing pipeline. **(1)** Heat maps of dLGN-p RGC Ca^2+^ responses to the to the chirp and the moving bar stimulus (n = 581, recorded with genetically encoded Ca^2+^-indicator GCaMP6f, responses sorted by experimental day). Each line represents the responses of an individual cell (activity color-coded, with warmer colors representing increased activity). **(2)** Histogram of cell quality indices (Qis) to chirp and bar stimulus. Only cells with Qi > 0.45 for the chirp and with Qi > 0.6 for the bar stimulus were considered for further processing. (Methods) **(3)** Heat maps of dLGN-p RGC responses passing (top, n = 251) or failing (bottom, n = 330) the Qi-thresholds, sorted by Qi. **(4)** Heat maps (like in **(1)**) of Ca^2+^ responses for all RGC clusters ("RGC-all" from ^3^, recorded with Ca^2+^-indicator OGB-1, n = 49 clusters). Responses are sorted by cluster index, with the height of a cluster 'block' representing the number of individual cells. **(5)** Heat maps of dLGN-p RGCs after the cluster assignment to RGC-all clusters, including trace deconvolution and trace correlation (Methods). The height of each cluster represents the number of included cells. **(6)** Distribution of dLGN-p RGCs per RGC-all cluster based on the cell cluster assignment. **(7)** Heat map of dLGN cell responses to chirp stimulus (responses sorted by experimental day). **(8)** Histograms of quality criteria, including, from top to bottom, quality index, Wilcoxon rank sum test for equal medians, cluster quality values, and STFO correlation values as computed by the linear regression analysis from two STFO stimuli played before and after the chirp. Cells had to be either above each of the top three quality criteria thresholds or the bottom one to pass into the analysis dataset. **(9)** Heat maps of dLGN cell responses that passed (top, n = 814) or failed (bottom, n = 1,376) the quality criteria, sorted by quality criteria values. **(10)** Schematic of the low-rank non-negative matrix factorization (NNMF), which was applied on the dLGN cell data from **(9)** Each neuron response is considered to be composed of particular temporal response components multiplied by neuron-specific weights. **(11)** OGB-1 Ca^2+^-kernel used for the convolution of dLGN spike traces into simulated OGB-1 Ca^2+^ traces (after down-sampling) with their characteristically slower Ca^2+^ response kinetics. This preprocessing step was applied so that the dLGN data better match the RGC data for the subsequent dLGN-model. **(12)** Heat map of dLGN responses after downsampling and convolution. **(13)** Illustration of the linear feedforward model of single dLGN cell responses from dLGN-p RGC cell types. The model is restricted to using only RGC clusters that received cell assignments in the dLGN-p RGC data set. It then predicts individual dLGN cell responses as a linear combination of weighted mean RGC-all cluster responses. Right of black arrow: Sample dLGN response (black) and its linear prediction (blue), and bar plot of contributing RGC-all cluster weights.

### 2. Deconvolution

**Supplementary Figure 2.**
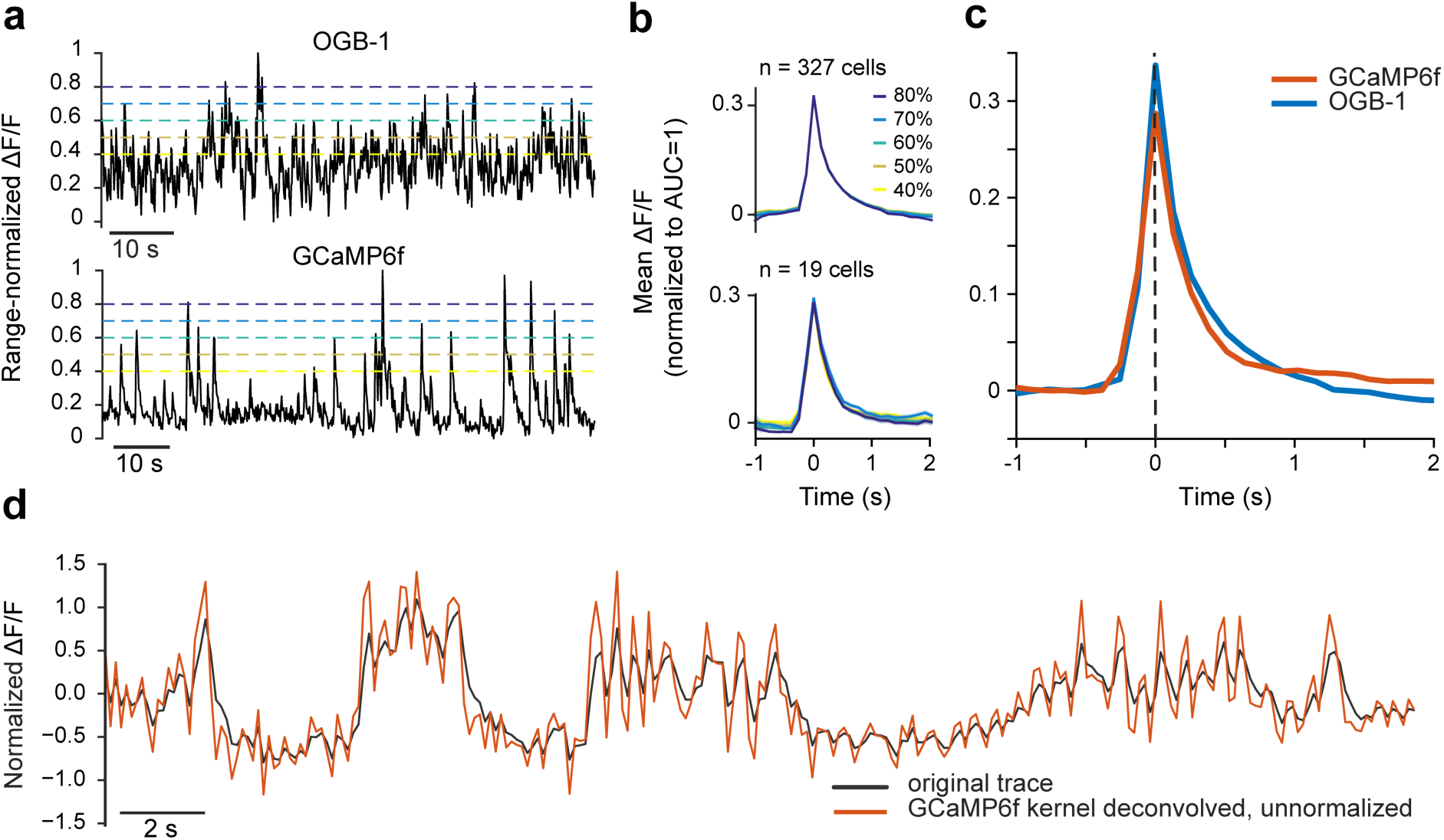
| Deconvolution. **a,** Raw traces to binary white noise stimulus with various calcium event thresholds (top: OGB-1; bottom: GCaMP6f). **b,** Extracted mean Ca^2+^ event kernels for OGB-1 (top) and GCaMP6f (bottom) for thresholds shown in (a). **c,** Superposition of OGB-1 and GCaMP6f kernels. **d,** Example GCaMP6f trace (in black), and deconvolution with the GCaMP6f kernel (orange).

### 3. Cluster assignment of dLGN-projecting RGCs

**Supplementary Figure 3.**
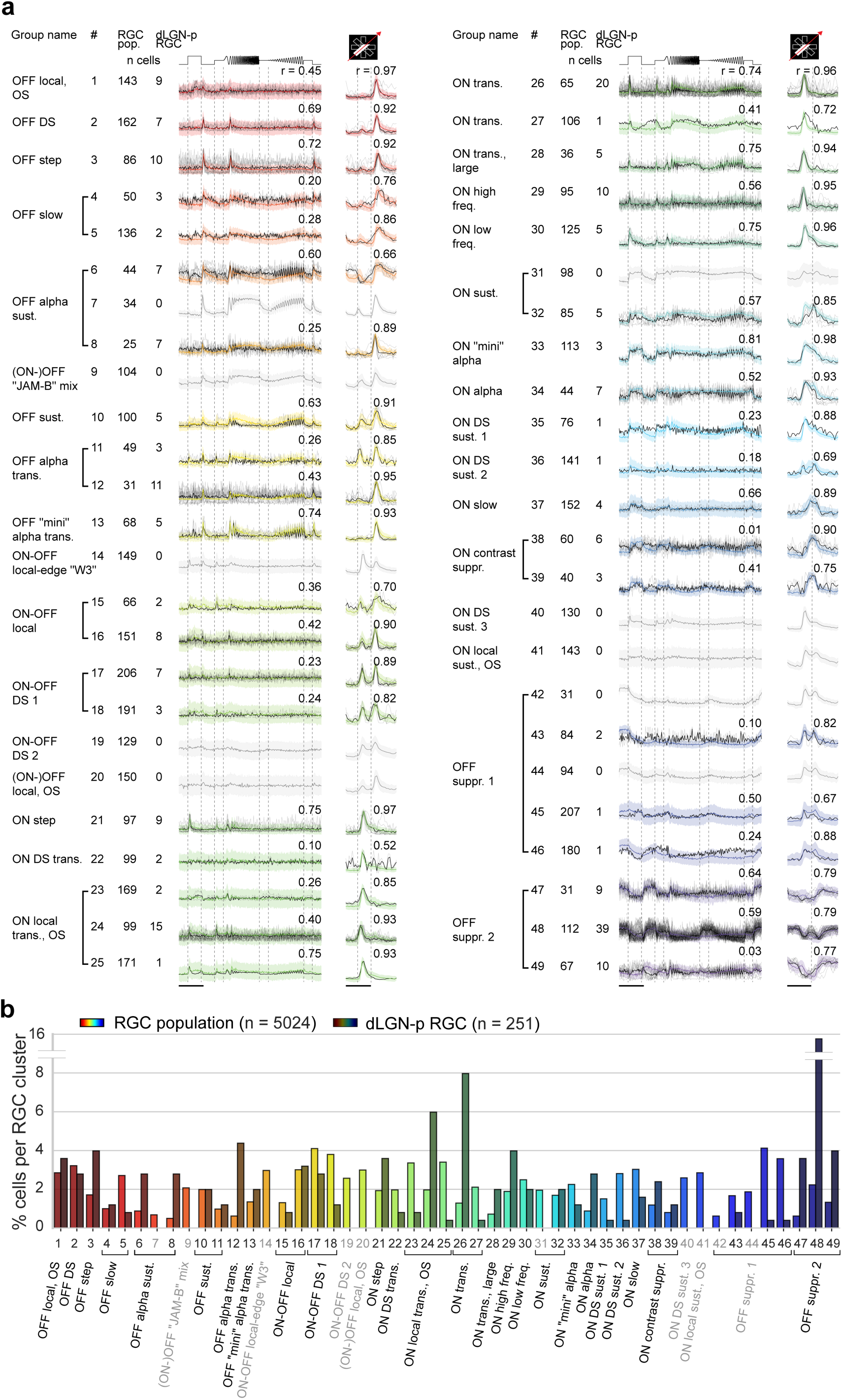
| dLGN-p RGC cluster assignments. **a,** All dLGN-p RGC cluster responses (grey: single RGCs; black: cluster mean), along with assigned RGC population cluster response mean (color) and SD (colored area). RGC population clusters that were not assigned any dLGN-p RGCs are greyed out. **b,** Percentage of cells per RGC cluster for dLGN-p RGCs (dark colors) and all RGCs obtained from ^3^ (saturated colors).

### 4. Responses of dLGN neurons to grating stimuli presented before and after the chirp stimulus

**Supplementary Figure 4.**
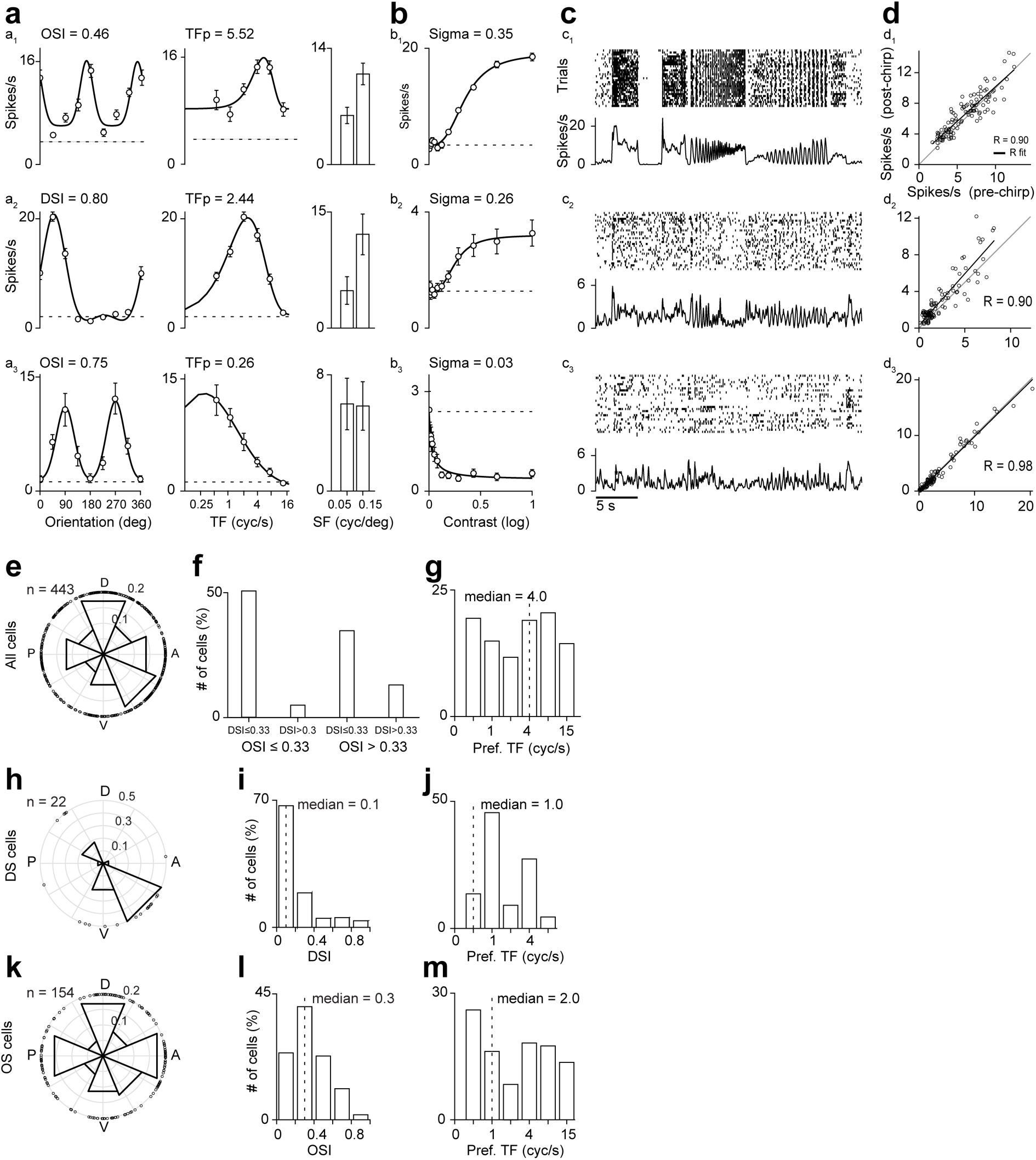
| dLGN responses to drifting gratings. **a-d,** Responses of three example dLGN cells to the full stimulus set. **a,** Mean firing rates and fitted tuning curves (red) for orientation, spatial frequency, and temporal frequency. **b,** Same as a, for contrast response functions. **c,** Responses to the chirp stimulus. **d,** Scatter plot of average firing rates across all conditions of the drifting grating stimuli presented before and after the chirp stimulus, used to determine the stability of the recorded cells. We chose R>0.65 as criterion to determine that a dLGN neuron was stable over time. **e-g,** Population data for direction and orientation selectivity: Distribution of preferred direction of motion for all responsive dLGN neurons (n = 443) (e); percentage of direction/orientation-selective cells, with a cut-off of 0.33 for DSI and OSI (f); distribution of preferred temporal frequencies (g). **h-j,** Same as (e-g), for DS cells (n = 22). **k-m,** Same as (e-g), for OS cells (n = 154).

### 5. Locomotion

**Supplementary Figure 5.**
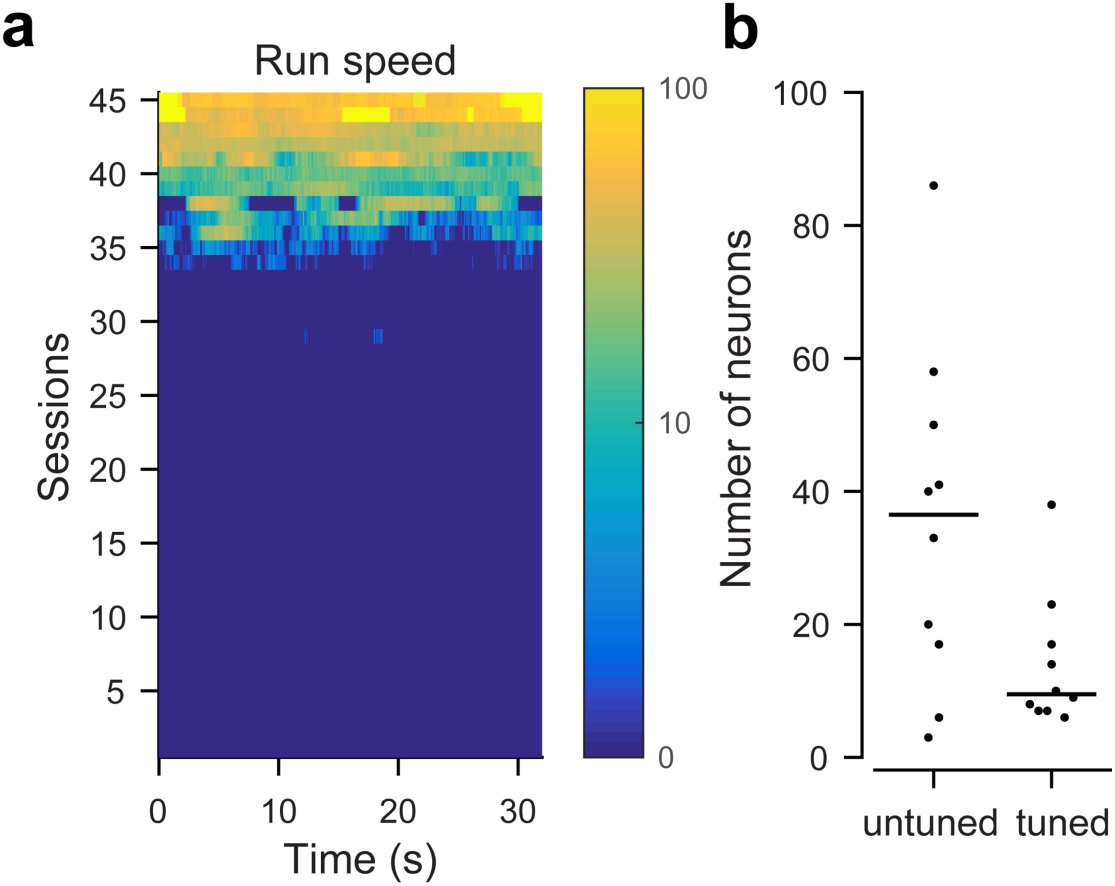
| Locomotion and run speed tuning of dLGN neurons. **a**, Average run speed (cm/s) per recording session as a function of time during the chirp stimulus. Locomotion speed was computed as the Euclidean norm of three perpendicular components of ball velocity ^90^. **b**, Number of neurons untuned and tuned for running speed per animal (n = 10 mice). We determined speed tuning as previously described in ^101^. In brief, speed traces were smoothed with a Gaussian filter (σ = 150 ms), re-sampled at 60 Hz, and binned such that each bin contained equal amounts of time (> 30 s). Unsmoothed neural responses were binned at 60 Hz. Neurons were considered speed modulated if the variance of mean responses across bins was greater than 99.9% of the variance of shuffled responses (p < 0.001).

### T1. Comparison of current data with the literature

**Supplementary Table 1.** Table showing putative correspondences between previously studied transgenic lines/markers for subgroups of RGCs following ^108^ and our functional RGC groups/clusters demonstrates an overall good match of our assignments of dLGN-p RGC groups with the literature. In case of JAM-B RGCs, there is a group labeled “ON-OFF JAM-B mix” in Baden et al. (2016), which is assigned no dLGN-p cells. However, it is uncertain whether this group really corresponds to JAM-B cells, which seem to be poorly responding to the stimulus set. For FSTL4, Drd4 and TRHR-cells, we chose ON-OFF DS1 as corresponding group in Baden et al. (2016), since the RGC types marked in these three lines are all ON-OFF DS and project to dLGN, as do cells in group 12. Correspondences in case of the CB2-and Opn4-lines to OFF alpha transient and ON alpha RGCs were based on the literature and the projection patterns matched well. We could not identify a match for Foxp2-RGCs. TYW3 cells were clearly identified in Baden et al. (2016) and were not among dLGN projecting RGCs. A subpopulation labeled W7A of TYW7 cells seems to correspond to OFF alpha sustained RGCs and does not project to dLGN. Interestingly, there was a cluster in the group labeled “OFF alpha sustained” in Baden et al. (2016) which was not assigned any dLGN-p cells. It is possible that the “OFF alpha sustained” group consists of heterogeneous RGC types. Alternatively, W7A cells are not “OFF alpha sustained” RGCs and all alpha RGCs project to dLGN. The three types of ON DS sustained in the Hoxd10 line do not project to the dLGN, and neither do the “ON DS sustained 1–3” groups in our data (although not for all of them the reduction in the representation is significant). Finally, the ON-OFF DS RGCs in the Hoxd10 line do not project to dLGN. Therefore, they most likely correspond to the ON-OFF DS 2 group of Baden et al., which is also not assigned any dLGN-p cells.

| transgenic line or marker | dLGN projections | reference | RGC name | group | cluster | dLGN-p in current dataset |
| --- | --- | --- | --- | --- | --- | --- |
| JAM-B-CreER | yes | 51,55 | JAM-B | 6? | 9? | no? |
| FSTL4-CreER | yes | 51,102 | ON-OFF DS 1 | 12 | 17, 18 | yes |
| Drd4-GFP | yes | 102,103 | ON-OFF DS 1 | 12 | 17, 18 | yes |
| TRHR-GFP | yes | 36 | ON-OFF DS 1 | 12 | 17, 18 | yes |
| CB2-GFP | yes | 104 | OFF alpha transient | 8 | 11, 12 | yes |
| Opn4-Cre | yes | 42,105,106 | ON alpha | 24 | 34 | yes |
| Foxp2 | yes | 107 | ? | ? | ? | ? |
| TYW3 | no | 50,51 | W3 | 10 | 14 | no |
| TYW7 | no | 51 | OFF alpha sust. | 5 | 7? | no? |
| Hoxd10-GFP | no | 52 | ON DS sust. 1 | 25 | 35 | no |
| Hoxd10-GFP, Spig1-GFP | no | 52-54 | ON DS sust. 2/3 | 26, 29 | 36, 40 | no |
| Hoxd10-GFP | no | 52 | ON-OFF DS 2 | 12, 13 | 19 | no |

## ACKNOWLEDGEMENTS

We thank G. Eske and M. Sotgia for excellent technical support and Alex H. Williams for discussion regarding NMF cross-validation. This research was funded by the German Research Foundation (DFG; SFB 1233 - Robust Vision: Inference Principles and Neural Mechanisms, TP 13; EXC 307, CIN; BE5601/4), the German Ministry for Science and Education (BMBF; FKZ 01GQ1002 and 01GQ1601), by the LMU Munich's Institutional Strategy LMUexcellent within the framework of the German Excellence Initiative (MRR), the SmartStart-program of the Bernstein Network for Computational Neuroscience funded by the Volkswagen Foundation (YB).

## Author contributions

TE, LB and PB designed the study; MRR and YB performed imaging experiments and pre-processing; MRR performed electrophysiological experiments; MRR and YB analysed the data; TE, LB and PB co-supervised the study; MRR, YB, TE, PB and LB wrote the manuscript.

## References

1. Euler, T., Haverkamp, S., Schubert, T. & Baden, T. Retinal bipolar cells: elementary building blocks of vision. Nat. Rev. Neurosci. 15, 507–19 (2014).

2. Masland, R. H. The Neuronal Organization of the Retina. Neuron 76, 266–280 (2012).

3. Baden, T. et al. The functional diversity of retinal ganglion cells in the mouse. Nature 529, 345–350 (2016).

4. Sanes, J. R. & Masland, R. H. The types of retinal ganglion cells: current status and implications for neuronal classification. Annu. Rev. Neurosci. 38, 221–46 (2015).

5. Martersteck, E. M. et al. Diverse Central Projection Patterns of Retinal Ganglion Cells. Cell Rep. 18, 2058–2072 (2017).

6. Morin, L. P. & Studholme, K. M. Retinofugal projections in the mouse. J. Comp. Neurol. 522, 3733–3753 (2014).

7. Huberman, A. D., & Niell, C. M. What can mice tell us about how vision works? Trends Neurosci. 34, 464–473 (2011).

8. Usrey, W. M., Reppas, J. B. & Reid, R. C. Specificity and strength of retinogeniculate connections. J. Neurophysiol. 82, 3527–3540 (1999).

9. Cleland, B. G., Dubin, M. W. & Levick, W. R. Simultaneous recording of input and output of lateral geniculate neurones. Nat. New Biol. 231, 191–192 (1971).

10. Hubel, D. H. & Wiesel, T. N. Integrative action in the cat’s lateral geniculate body. J. Physiol. 155, 385–398 (1961).

11. Kaplan, E., Purpura, K. & Shapley, R. M. Contrast affects the transmission of visual information through the mammalian lateral geniculate nucleus. J. Physiol. 391, 267–288 (1987).

12. Sincich, L. C., Adams, D. L., Economides, J. R. & Horton, J. C. Transmission of spike trains at the retinogeniculate synapse. J. Neurosci. 27, 2683–2692 (2007).

13. Hamos, J. E., Van Horn, S. C., Raczkowski, D. & Sherman, S. M. Synaptic circuits involving an individual retinogeniculate axon in the cat. J. Comp. Neurol. 259, 165–192 (1987).

14. Mastronarde, D. N. Two classes of single-input X-cells in cat lateral geniculate nucleus. Retinal inputs and the generation of receptive-field properties. J. Neurophysiol. 57, 381–413 (1987).

15. Mastronarde, D. N. Nonlagged relay cells and interneurons in the cat lateral geniculate nucleus: receptive-field properties and retinal inputs. Vis. Neurosci. 8, 407–441 (1992).

16. Dacey, D. M., Peterson, B. B., Robinson, F. R. & Gamlin, P. D. Fireworks in the primate retina: in vitro photodynamics reveals diverse LGN-projecting ganglion cell types. Neuron 37, 15–27 (2003).

17. Rompani, S. B. et al. Different Modes of Visual Integration in the Lateral Geniculate Nucleus Revealed by Single-Cell-Initiated Transsynaptic Tracing. Neuron 93, 767–-776.e6 (2017).

18. Hammer, S., Monavarfeshani, A., Lemon, T., Su, J. & Fox, M. A. Multiple Retinal Axons Converge onto Relay Cells in the Adult Mouse Thalamus. Cell Rep. 12, 1575–1583 (2015).

19. Morgan, J. L., Berger, D. R., Wetzel, A. W. & Lichtman, J. W. The Fuzzy Logic of Network Connectivity in Mouse Visual Thalamus. Cell 165, 192–206 (2016).

20. Baden, T. et al. The functional diversity of mouse retinal ganglion cells. Nature 529, 1–21 (2016).

21. Rompani, S. B. et al. Different Modes of Visual Integration in the Lateral Geniculate Nucleus Revealed by Single-Cell-Initiated Transsynaptic Tracing. 767–776 (2017). doi:10.1016/j.neuron.2017.01.028

22. Denman, D. J. & Contreras, D. On Parallel Streams through the Mouse Dorsal Lateral Geniculate Nucleus. Front. Neural Circuits 10, 20 (2016).

23. Grubb, M. S. & Thompson, I. D. Quantitative characterization of visual response properties in the mouse dorsal lateral geniculate nucleus. J. Neurophysiol. 90, 3594–3607 (2003).

24. Grubb, M. S. & Thompson, I. D. Quantitative characterization of visual response properties in the mouse dorsal lateral geniculate nucleus. J. Neurophysiol. 90, 3594–3607 (2003).

25. Marshel, J. H., Kaye, A. P., Nauhaus, I. & Callaway, E. M. Anterior-Posterior Direction Opponency in the Superficial Mouse Lateral Geniculate Nucleus. Neuron 76, 713–720 (2012).

26. Piscopo, D. M., El-Danaf, R. N., Huberman, A. D. & Niell, C. M. Diverse Visual Features Encoded in Mouse Lateral Geniculate Nucleus. J. Neurosci. 33, 4642–4656 (2013).

27. Zhao, X., Chen, H., Liu, X. & Cang, J. Orientation-selective responses in the mouse lateral geniculate nucleus. J. Neurosci. 33, 12751–12763 (2013).

28. Neve, R. L. Overview of gene delivery into cells using HSV-1-based vectors. Curr. Protoc. Neurosci. Chapter 4, Unit 4.12 (2012).

29. Chen, T.-W. et al. Ultrasensitive fluorescent proteins for imaging neuronal activity. Nature 499, 295–300 (2013).

30. Madisen, L. et al. Transgenic mice for intersectional targeting of neural sensors and effectors with high specificity and performance. Neuron 85, 942–958 (2015).

31. Antinone, S. E. & Smith, G. A. Retrograde axon transport of herpes simplex virus and pseudorabies virus: a live-cell comparative analysis. J. Virol. 84, 1504–1512 (2010).

32. McGavern, D. B. & Kang, S. S. Illuminating viral infections in the nervous system. Nat. Rev. Immunol. 11, 318–329 (2011).

33. Guillery, R. W. & Sherman, S. M. Thalamic relay functions and their role in corticocortical communication: generalizations from the visual system. Neuron 33, 163–175 (2002).

34. Harting, J. K., Huerta, M. F., Hashikawa, T. & van Lieshout, D. P. Projection of the mammalian superior colliculus upon the dorsal lateral geniculate nucleus: organization of tectogeniculate pathways in nineteen species. J. Comp. Neurol. 304, 275–306 (1991).

35. Ellis, E. M., Gauvain, G., Sivyer, B. & Murphy, G. J. Shared and distinct retinal input to the mouse superior colliculus and dorsal lateral geniculate nucleus. J. Neurophysiol. 116, 602–610 (2016).

36. Rivlin-Etzion, M. et al. Transgenic mice reveal unexpected diversity of on-off direction-selective retinal ganglion cell subtypes and brain structures involved in motion processing. J. Neurosci. 31, 8760–8769 (2011).

37. Lee, D. D. & Seung, H. S. Learning the parts of objects by non-negative matrix factorization. Nature 401, 788–791 (1999).

38. Bro, R., Kjeldahl, K., Smilde, A. K. & Kiers, H. A. L. Cross-validation of component models: A critical look at current methods. Anal. Bioanal. Chem. 390, 1241–1251 (2008).

39. van Wyk, M., Wässle, H. & Taylor, W. R. Receptive field properties of ON- and OFF-ganglion cells in the mouse retina. Vis. Neurosci. 26, 297–308 (2009).

40. Krieger, B., Qiao, M., Rousso, D. L., Sanes, J. R. & Meister, M. Four alpha ganglion cell types in mouse retina: Function, structure, and molecular signatures. PLoS One 12, 1–21 (2017).

41. Brown, T. M. et al. Melanopsin contributions to irradiance coding in the thalamo-cortical visual system. PLoS Biol. 8, e1000558 (2010).

42. Ecker, J. L. et al. Melanopsin-expressing retinal ganglion-cell photoreceptors: Cellular diversity and role in pattern vision. Neuron 67, 49–60 (2010).

43. Hattar, S. et al. Central projections of melanopsin-expressing retinal ganglion cells in the mouse. J. Comp. Neurol. 497, 326–349 (2006).

44. Niell, C. M. & Stryker, M. P. Modulation of Visual Responses by Behavioral State in Mouse Visual Cortex. Neuron 65, 472–479 (2010).

45. Tien, N.-W., Pearson, J. T., Heller, C. R., Demas, J. & Kerschensteiner, D. Genetically Identified Suppressed-by-Contrast Retinal Ganglion Cells Reliably Signal {Self-Generated} Visual Stimuli. J. Neurosci. 35, 10815–10820 (2015).

46. Levick, W. R. Receptive fields and trigger features of ganglion cells in the visual streak of the rabbits retina. J. Physiol. 188, 285–307 (1967).

47. Masland, R. H. & Martin, P. R. The unsolved mystery of vision. Curr. Biol. 17, R577–-82 (2007).

48. Sivyer, B., Taylor, W. R. & Vaney, D. I. Uniformity detector retinal ganglion cells fire complex spikes and receive only light-evoked inhibition. Proc. Natl. Acad. Sci. U. S. A. 107, 5628–5633 (2010).

49. Troy, J. B., Einstein, G., Schuurmans, R. P., Robson, J. G. & Enroth-Cugell, C. Responses to sinusoidal gratings of two types of very nonlinear retinal ganglion cells of cat. Vis. Neurosci. 3, 213–223 (1989).

50. Zhang, Y., Kim, I.-J., Sanes, J. R. & Meister, M. The most numerous ganglion cell type of the mouse retina is a selective feature detector. Proc. Natl. Acad. Sci. U. S. A. 109, E2391–-8 (2012).

51. Kim, I.-J., Zhang, Y., Meister, M. & Sanes, J. R. Laminar restriction of retinal ganglion cell dendrites and axons: subtype-specific developmental patterns revealed with transgenic markers. J. Neurosci. 30, 1452–1462 (2010).

52. Dhande, O. S. et al. Genetic dissection of retinal inputs to brainstem nuclei controlling image stabilization. J. Neurosci. 33, 17797–17813 (2013).

53. Yonehara, K. et al. Expression of SPIG1 reveals development of a retinal ganglion cell subtype projecting to the medial terminal nucleus in the mouse. PLoS One 3, e1533 (2008).

54. Yonehara, K. et al. Identification of retinal ganglion cells and their projections involved in central transmission of information about upward and downward image motion. PLoS One 4, e4320 (2009).

55. Kim, I.-J., Zhang, Y., Yamagata, M., Meister, M. & Sanes, J. R. Molecular identification of a retinal cell type that responds to upward motion. Nature 452, 478–482 (2008).

56. Pang, J.-J., Gao, F. & Wu, S. M. Light-evoked excitatory and inhibitory synaptic inputs to ON and OFF alpha ganglion cells in the mouse retina. J. Neurosci. 23, 6063–6073 (2003).

57. Kerschensteiner, D. & Guido, W. Organization of the dorsal lateral geniculate nucleus in the mouse. Vis. Neurosci. 34, E008 (2017).

58. Howarth, M., Walmsley, L. & Brown, T. M. Binocular integration in the mouse lateral geniculate nuclei. Curr. Biol. 24, 1241–1247 (2014).

59. Cruz-Martín, A. et al. A dedicated circuit links direction-selective retinal ganglion cells to the primary visual cortex. Nature 507, 358–61 (2014).

60. Scholl, B., Tan, A. Y. Y., Corey, J. & Priebe, N. J. Emergence of Orientation Selectivity in the Mammalian Visual Pathway. J. Neurosci. 33, 10616–10624 (2013).

61. Hei, X. et al. Directional selective neurons in the awake LGN: response properties and modulation by brain state. J. Neurophysiol. 112, 362–373 (2014).

62. Zeater, N., Cheong, S. K., Solomon, S. G., Dreher, B. & Martin, P. R. Binocular Visual Responses in the Primate Lateral Geniculate Nucleus. Curr. Biol. 25, 3190–3195 (2015).

63. White, a J., Solomon, S. G. & Martin, P. R. Spatial properties of koniocellular cells in the lateral geniculate nucleus of the marmoset Callithrix jacchus. J. Physiol. 533, 519–535 (2001).

64. Cheong, S. K., Tailby, C., Solomon, S. G. & Martin, P. R. Cortical-Like Receptive Fields in the Lateral Geniculate Nucleus of Marmoset Monkeys. J. Neurosci. 33, 6864–6876 (2013).

65. Chen, C. & Regehr, W. G. Developmental remodeling of the retinogeniculate synapse. Neuron 28, 955–966 (2000).

66. Weyand, T. G. The multifunctional lateral geniculate nucleus. Rev. Neurosci. 27, 135–157 (2016).

67. Jaubert-Miazza, L. et al. Structural and functional composition of the developing retinogeniculate pathway in the mouse. Vis. Neurosci. 22, 661–676 (2005).

68. Litvina, E. Y. & Chen, C. Functional Convergence at the Retinogeniculate Synapse. Neuron 96, 330–-338.e5 (2017).

69. Reid, R. C. & Usrey, W. M. Functional connectivity in the pathway from retina to striate cortex. Vis. Neurosci. 1, 673–679 (2004).

70. Ziburkus, J. & Guido, W. Loss of binocular responses and reduced retinal convergence during the period of retinogeniculate axon segregation. J. Neurophysiol. 96, 2775–2784 (2006).

71. Morgan, J. L., Berger, D. R., Wetzel, A. W. & Lichtman, J. W. The Fuzzy Logic of Network Connectivity in Mouse Visual Thalamus. Cell 165, 192–206 (2016).

72. Hammer, S., Monavarfeshani, A., Lemon, T., Su, J. & Fox, M. A. Multiple Retinal Axons Converge onto Relay Cells in the Adult Mouse Thalamus. Cell Rep. 12, 1575–1583 (2015).

73. Chen, C., Bickford, M. E. & Hirsch, J. A. Untangling the Web between Eye and Brain. Cell 165, 20–21 (2016).

74. Hong, Y. K. et al. Refinement of the Retinogeniculate Synapse by Bouton Clustering. Neuron 84, 332–339 (2014).

75. Rathbun, D. L., Alitto, H. J., Warland, D. K. & Usrey, W. M. Stimulus Contrast and Retinogeniculate Signal Processing. Front. Neural Circuits 10, (2016).

76. Martinez, L. M., Molano-Mazón, M., Wang, X., Sommer, F. T. & Hirsch, J. A. Statistical wiring of thalamic receptive fields optimizes spatial sampling of the retinal image. Neuron 81, 943–956 (2014)

77. Sillito, A. M., Cudeiro, J. & Jones, H. E. Always returning: feedback and sensory processing in visual cortex and thalamus. Trends Neurosci. 29, 307–316 (2006).

78. Fitzpatrick, D., Diamond, I. T. & Raczkowski, D. Cholinergic and monoaminergic innervation of the cat’s thalamus: Comparison of the lateral geniculate nucleus with other principal sensory nuclei. J. Comp. Neurol. 288, 647–675 (1989).

79. Antal, M., Acuna-Goycolea, C., Todd Pressler, R., Blitz, D. M. & Regehr, W. G. Cholinergic activation of M2 receptors leads to context-dependent modulation of feedforward inhibition in the visual thalamus. PLoS Biol. 8, (2010).

80. Hirsch, J. A., Wang, X., Sommer, F. T. & Martinez, L. M. How Inhibitory Circuits in the Thalamus Serve Vision. Annu. Rev. Neurosci. 38, (2015).

81. Lesica, N. A. et al. Dynamic encoding of natural luminance sequences by LGN bursts. PLoS Biol. 4, 1201–1212 (2006).

82. Erisken, S. et al. Effects of locomotion extend throughout the mouse early visual system. Curr. Biol. 24, 2899–2907 (2014).

83. Bezdudnaya, T. et al. Thalamic burst mode and inattention in the awake LGNd. Neuron 49, 421–432 (2006).

84. Bouabe, H. & Okkenhaug, K. Gene targeting in mice: A review. Methods in Molecular Biology 1064, 315–336 (2013).

85. Vaiceliunaite, A., Erisken, S., Franzen, F., Katzner, S. & Busse, L. Spatial integration in mouse primary visual cortex. J. Neurophysiol. 110, 964–72 (2013).

86. Euler, T. et al. Eyecup scope-optical recordings of light stimulus-evoked fluorescence signals in the retina. Pflugers Arch. Eur. J. Physiol. 457, 1393–1414 (2009).

87. Baden, T. et al. A tale of two retinal domains: near-optimal sampling of achromatic contrasts in natural scenes through asymmetric photoreceptor distribution. Neuron 80, 1206–1217 (2013).

88. Dorostkar, M. M., Dreosti, E., Odermatt, B. & Lagnado, L. Computational processing of optical measurements of neuronal and synaptic activity in networks. J. Neurosci. Methods 188, 141–150 (2010).

89. Ecker, A. S. et al. State dependence of noise correlations in macaque primary visual cortex. Neuron 82, 235–248 (2014).

90. Dombeck, D. A., Khabbaz, A. N., Collman, F., Adelman, T. L. & Tank, D. W. Imaging large-scale neural activity with cellular resolution in awake, mobile mice. Neuron 56, 43–57 (2007).

91. Yatsenko, D. et al. DataJoint: managing big scientific data using MATLAB or Python. (2015).

92. Hazan, L., Zugaro, M. & Buzsáki, G. Klusters, NeuroScope, NDManager: a free software suite for neurophysiological data processing and visualization. J. Neurosci. Methods 155, 207–216 (2006).

93. Quiroga, R. Q., Nadasdy, Z. & Ben-Shaul, Y. Unsupervised spike detection and sorting with wavelets and superparamagnetic clustering. Neural Comput. 16, 1661–1687 (2004).

94. Henze, D. A. et al. Intracellular features predicted by extracellular recordings in the hippocampus in vivo. J. Neurophysiol. 84, 390–400 (2000).

95. Liu, B.-H. et al. Intervening inhibition underlies simple-cell receptive field structure in visual cortex. Nat. Neurosci. 13, 89–96 (2010).

96. Albrecht, D. G. & Hamilton, D. B. Striate cortex of monkey and cat: contrast response function. J. Neurophysiol. 48, 217–237 (1982).

97. Katzner, S., Busse, L. & Carandini, M. GABAA inhibition controls response gain in visual cortex. J. Neurosci. 31, 5931–5941 (2011).

98. Niell, C. M., & Stryker, M. P. Highly selective receptive fields in mouse visual cortex. J. Neurosci. 28, 7520–7536 (2008).

99. Wold, S. Cross-Validatory Estimation of the Number of Components in Factor and Principal Components Models. Technometrics 20, 397–405 (1978).

100. Ward, J. H. Hierarchical Grouping to Optimize an Objective Function. J. Am. Stat. Assoc. 58, 236–244 (1963).

101. Saleem, A. B., Ayaz, A., Jeffery, K. J., Harris, K. D. & Carandini, M. Integration of visual motion and locomotion in mouse visual cortex. Nat. Neurosci. 16, 1864–1869 (2013).

102. Kay, J. N. et al. Retinal ganglion cells with distinct directional preferences differ in molecular identity, structure, and central projections. J. Neurosci. 31, 7753–7762 (2011).

103. Huberman, A. D. et al. Genetic Identification of an On-Off Direction-Selective Retinal Ganglion Cell Subtype Reveals a Layer-Specific Subcortical Map of Posterior Motion. Neuron 62, 327–334 (2009).

104. Huberman, A. D. et al. Architecture and Activity-Mediated Refinement of Axonal Projections from a Mosaic of Genetically Identified Retinal Ganglion Cells. Neuron 59, 425–438 (2008).

105. Estevez, M. E. et al. Form and function of the M4 cell, an intrinsically photosensitive retinal ganglion cell type contributing to geniculocortical vision. J. Neurosci. 32, 13608–20 (2012).

106. Schmidt, T. M. et al. A role for melanopsin in alpha retinal ganglion cells and contrast detection. Neuron 82, 781–788 (2014).

107. Rousso, D. L. et al. Two Pairs of ON and OFF Retinal Ganglion Cells Are Defined by Intersectional Patterns of Transcription Factor Expression. Cell Rep. 15, 1930–1944 (2016).

108. Dhande, O. S., Stafford, B. K., Lim, J.-H. A. & Huberman, A. D. Contributions of Retinal Ganglion Cells to Subcortical Visual Processing and Behaviors. Annu. Rev. Vis. Sci. 1, 291–328 (2015).

